# Brain responses vary in duration - modeling strategies and challenges

**DOI:** 10.1101/2024.12.05.626938

**Authors:** René Skukies, Judith Schepers, Benedikt Ehinger

## Abstract

Typically, event-related brain responses are calculated invariant to the underlying event duration, even in cases where event durations observably vary: with reaction times, fixation durations, word lengths, or varying stimulus durations. Additionally, an often co-occurring consequence of differing event durations is a variable overlap of the responses to subsequent events. While the problem of overlap e.g. in fMRI and EEG is successfully addressed using linear overlap correction, it is unclear whether overlap correction and duration covariate modeling can be jointly used, as both are dependent on the same inter-event-distance variability. Here, we first show that failing to explicitly account for event durations can lead to spurious results and thus are important to consider. Next, we propose and compare several methods based on multiple regression to explicitly account for stimulus durations. Using simulations, we find that non-linear spline regression of the duration effect outperforms other candidate approaches. Finally, we show that non-linear event duration modeling is compatible with linear overlap correction in time, making mass-univariate models combining non-linear spline regression of duration and linear overlap correction a flexible and appropriate tool to model overlapping brain signals. This allows us to reconcile the analysis of brain responses to stimuli in situations where durations differ between conditions, e.g. different reaction times, different stimulus durations, or fixation-related potentials with different fixation durations. While in this paper we focus on EEG analyses, we additionally show that our findings generalize to fMRI BOLD-responses, and argue that they should generalize to other overlapping signals such as LFPs, or pupil dilation responses.

## 1. Introduction

### 1.1. Event-Related Potentials

Neural activity is rarely interpretable without removing unwanted noise through averaging. Such averaging is most commonly performed time-point by time-point over multiple trial repetitions relative to an event marker. Under the assumption of uncorrelated noise, averaging will, in the limit, remove all interferences which are inconsistent across trials, recovering the “true” underlying event-related signal.

In human EEG, the result of such averaging is known as the event-related potential (ERP) and has been studied for more than 80 years (Davis, 1936). While in this article we primarily focus on such ERPs, we additionally demonstrate that our findings generalize to other event-related time series on the example of an fMRI dataset.

A helpful example to illustrate the process of calculating an ERP can be found in its application to a classical P300 experiment, commonly known as an active oddball experiment. Here, subjects respond whether they see a rare “target” stimulus, or a frequent “distractor” stimulus (Figure 2.A). Typically, the signal of interest is the time-locked activity to the stimulus and, in some analyses, to the button press as well (Jung et al., 1999). After averaging several trials, a difference in the P300 between target and distractor stimuli manifests at around 350ms after the stimulus onset (Kappenman et al., 2021; Luck, 2014). While single-trial analyses exist, particularly in brain-computer interfaces, the averaging step has been pivotal to the development of ERP research.

### 1.2. Event Durations as Confounding Factor

This classical averaging step, however, cannot account for trial-wise (confounding) influences. For instance, in averaging we typically assume that a “stationary” ERP exists on every single trial, but is contaminated with noise. If we suspect that other effects can affect the overall shape of the ERP (or another event related measure of interest) in space and/ or time, then we can no longer use simple averaging as our analysis method (see Figure 1 for simulations of exemplary duration effects in pupil dilation (A), fMRI-BOLD (B), and fixation-related potentials(C)). Examples of this problem and how to address it have been discussed and spearheaded by Pernet & Rousselet in their LIMO approach (Pernet et al., 2011), with the rERP approach (Smith & Kutas, 2015a) and also in our own work for low-level stimulus confounds (Ehinger & Dimigen, 2019) and for eye-movement attributes (Dimigen & Ehinger, 2021).

**Figure 1.**
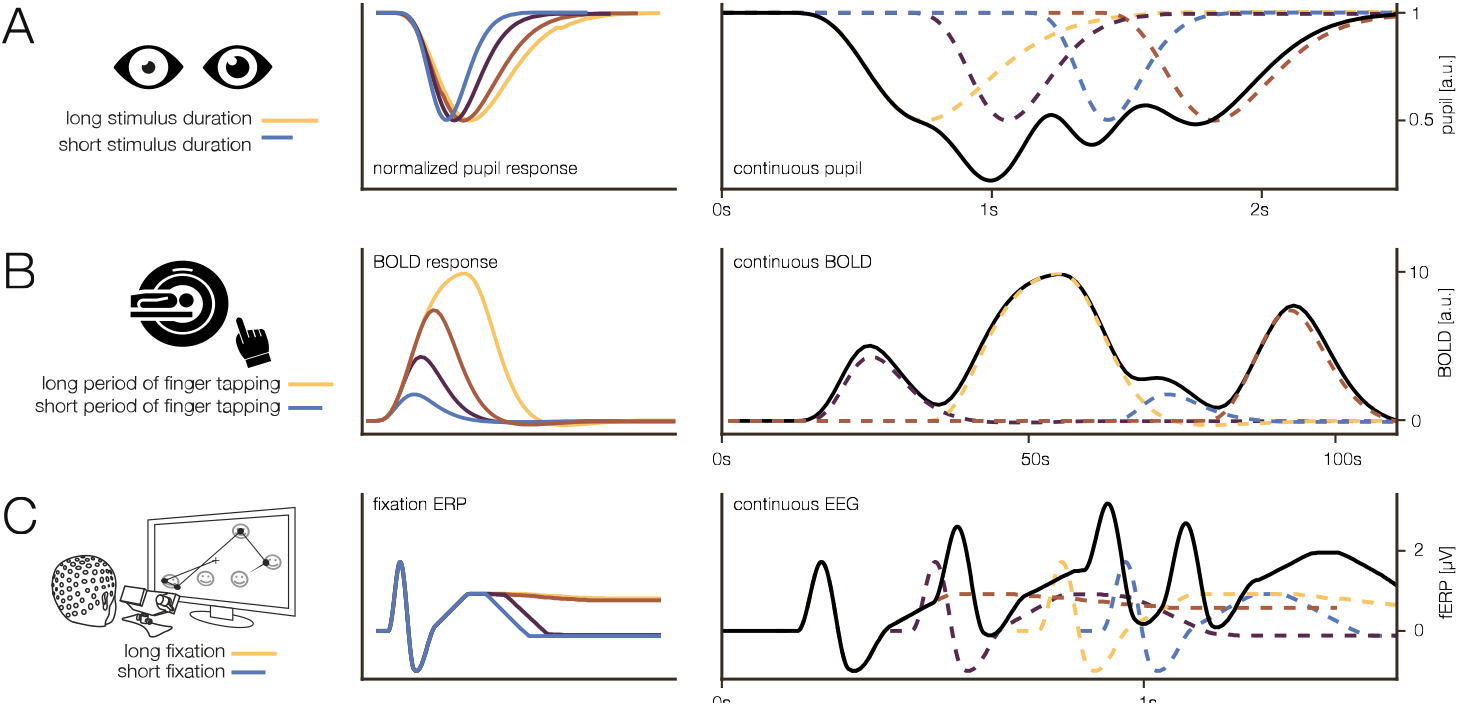
Exemplary duration effects in different modalities. Simulations following published effects (Pupil: Snowden et al. (2016), BOLD: Glover (1999), FRP: this paper). A) normalized pupil response to varying stimulus durations. B) BOLD response to different block durations of finger tapping. C) FRP responses to fixations of varying durations.

**Figure 2.**
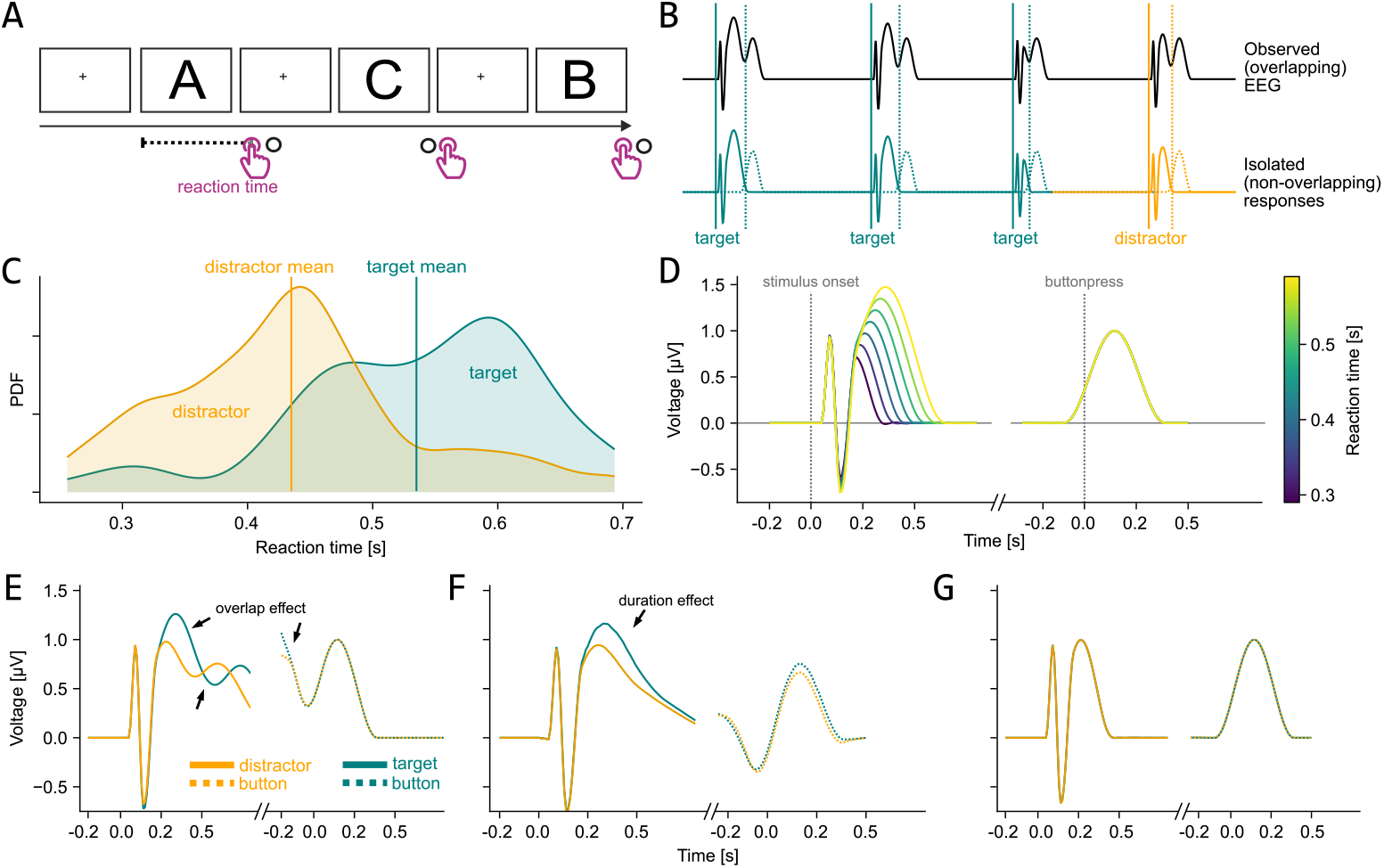
Potential influence of reaction time in an oddball task. (A) Experimental sequence of the classical active oddball task; Because the next stimulus is shown only after the response of the participants, their reaction time controls the duration of a trial. (B) Observed EEG during the task (top) and underlying single-event responses (bottom; solid lines = stimulus onset/ERP, dashed line = response onset/ERP). (C) Distribution of reaction times between conditions of a single subject in an active oddball task from the ERP core dataset (Kappenman et al., 2021). (D) Simulated isolated response to stimulus and response, including the effect of reaction time on the stimulus response. (E) Results from a classical average ERP analysis. Many artefactual differences are visible. (F) Average ERP after overlap correction. (G) ERP after overlap correction and covariate control through general additive modeling, showing that a condition effect is only attributed to a difference in reaction time distribution (E-G are results from simulated data). For a similar figure showcasing fixation-related potentials and the effect of fixation duration, see Figure 2 in Dimigen & Ehinger (2021) and for face-related activity see Ehinger & Dimigen (2019).

Varying event durations likely reflect one such trial-wise influence (Figure 2.D), manifested through external stimulation (e.g. how long a stimulus is presented in the experiment), as well as internal processes (e.g. how long a stimulus is processed by the brain, often indirectly measured via reaction time).

Especially the latter is a strong candidate for potential confounds, as stimulus processing times can be decoded from many brain areas using fMRI (Nakuci et al., 2024), but more importantly, typically differ between conditions (Gilbert & Sigman, 2007; Lange & Röder, 2006), and populations (Der & Deary, 2006) (see Figure 2.C for different reaction times between conditions in an oddball experiment). Processing times also naturally differ for fixation durations, e.g. in unrestricted viewing (Nuthmann, 2017), or for certain stimuli, e.g. faces, which are typically fixated longer than other objects (Gert et al., 2022).

Indeed, previous studies by Wang et al. (2018), Hassall et al. (2022), Yarkoni et al. (2009), Groen et al. (2022), Brands et al. (2024) & Mumford et al. (2023) have shown the importance to regard duration as a potential confounder in single-neuron activity, in EEG, iEEG and also fMRI. Wang et al. (2018) recorded single-neuron activity from the medial frontal cortex in non-human primates, which performed a task to produce intervals of motor activity (i.e. a hand movement towards a stimulus) of different lengths. Through this, they showed that neural activity temporally scaled with interval length. Building on this research, Hassall et al. (2022) extended the idea of temporally scaled signals to human cognition. By using a general linear model containing fixed- and variable-duration regressors, they successfully unmixed fixed- and scaled-time components in human EEG data. Furthermore, Sun et al. (2024) recently expressed concerns about reaction time, a prominent example of event duration, as an important confounding variable in response-locked ERPs. Adding to this, Yarkoni et al. (2009) and Mumford et al. (2023) emphasized the importance of considering reaction time in fMRI time series modeling, showing that a reaction time effect can be found over a wide area of brain regions in multiple tasks. Further, modelling duration effects in EEG could allow to include the within-subject effect in subsequent fMRI analyses as proposed by Philiastides & Sajda (2007). In their work, they identified components in EEG data of participants doing a decision-making task. Using this information as regressors in the analysis of fMRI data, they received better estimate of the spatiotemporal effect of decision making. In summary, these studies highlight the increasing importance of considering event duration as a crucial factor in neural and cognitive research. And indeed, researchers have explicitly started to include event duration as a predictor in their models (Nikolaev et al., 2023).

As mentioned above, one promising solution to address trial-wise influences through event duration is the regression ERP (rERP) framework (Smith & Kutas, 2015b). Here, a mass univariate multiple regression is applied to all time-points of an epoch around the event of interest (Frömer et al., 2018; Hauk et al., 2006; Pernet et al., 2011; Smith & Kutas, 2015a). This flexible framework allows us to incorporate covariates varying over trials, like reaction time, fixation duration, or stimulus duration, and subsequently statistically adjust for covariate imbalances between conditions (e.g., a faster reaction time towards the distractors than the targets in the P300 experiment). The rERP approach further allows employing additive models that break the shackles of linear covariate modeling and allow for arbitrary smooth dependencies, which seem appropriate for complex neural activity as expected by duration effects. In the last few years, the workflow using these approaches greatly improved, as new, specialized, and user-friendly toolboxes e.g. LIMO (Pernet et al., 2011), or Unfold (Dimigen & Ehinger, 2021; Ehinger & Dimigen, 2019) were developed.

### 1.3. Overlapping Events

Having a method to tackle the duration problem still leaves us with the issue of temporal overlap between adjacent events, however. For instance, during an active oddball task, participants are asked to react with the press of a button to a row of different stimuli. Importantly, one rarely occurring stimulus is the designated target (i.e. the oddball; frequently occurring “non-oddball” stimuli are referred to as distractors) where participants have to react by use of a different button (Figure 2.A). Because participants’ responses to distractors occurs (for distractor targets) on average at about 430ms after stimulus onset, while a typical stimulus ERP lasts for longer than 600ms, the ERPs of stimuli and response will necessarily overlap with each other (Figure 2.B-C). Prior research has shown that such overlap, can be addressed within the same regression framework that is used to solve the covariate problem, by using linear overlap correction (Ehinger & Dimigen, 2019; Smith & Kutas, 2015b). This approach has been successfully applied to several experiments, with application to free viewing tasks (e.g. Coco et al., 2020; Gert et al., 2022; Nikolaev et al., 2023; Welke & Vessel, 2022), language modeling (Momenian et al., 2024), auditory modeling (Skerritt-Davis & Elhilali, 2018) and many other fields.

Yet, this leaves us with a conundrum: the linear overlap correction works due to the varying-event timing, but the varying-event timing also relates to the duration effect. Therefore, it is not at all clear whether these two factors, overlap and varying event duration, can, should, or even need to be modeled concurrently.

Here, we propose to incorporate event duration as a covariate as well as linear overlap correction into one coherent model to estimate the time course of an ERP. If possible, we could disentangle the influence of both factors and get a better estimate of the true underlying ERP (see Figure 2.E-G for simulated examples of results where: neither overlap nor a duration effect is accounted for (E); only overlap is corrected (F); and both the duration effect and overlap is corrected (G)). We further propose to use non-linear spline regression to model the duration effect in a smoother and more flexible way than would be possible with linear regression.

To show the validity of our approach, we use systematic simulations and compare spline regression with multiple alternative ways of modeling event duration. We further explore potential interactions with overlapping signals. Lastly, we demonstrate the implications of our approach by applying it to a real free-viewing dataset as well as to an fMRI stroop task.

## 2. Simulations

### 2.1. Approach

We sought to explore whether it is possible to model both event duration and overlap, two co-dependent factors, in one combined model. However, a problem with real-world EEG data is that we have no access to the true underlying ERP. Thus, we first turned towards simulations for which we can generate known “ground truths”, which vary with a continuous factor of duration, and which can be combined into one overlapping continuous signal by specifying the inter-event-onsets. Subsequently, we can apply our proposed analysis method and directly compare the results with the ground truth. All simulations were done in Matlab using custom scripts, which are publicly available at: https://github.com/s-ccs/unfold_duration

As “proto-ERPs” (i.e. ground truth) representing our duration-modulated event signal, we used a hanning window, varying the overall ERP shape in three different ways: stretching the hanning window along both axes, i.e. in time and amplitude (Figure 3.A) left panel; termed “scaled hanning”); stretching the overall ERP shape along the x-axis by the duration effect (Figure 3.A) middle panel; termed “hanning”); and stretching the later half along the x-axis by the duration effect, i.e. in time (Figure 3.A right panel; termed “half hanning”). We selected this set of shapes as we believe that they capture a wide range of the parameter space of a changing shape, and they would be better explainable than more complex shapes. Additionally, other, more complex shapes, e.g. based on adding duration-independent positive and negative peaks (like in a P1-N1-P3 ERP) did not show any qualitative difference in results.

**Figure 3.**
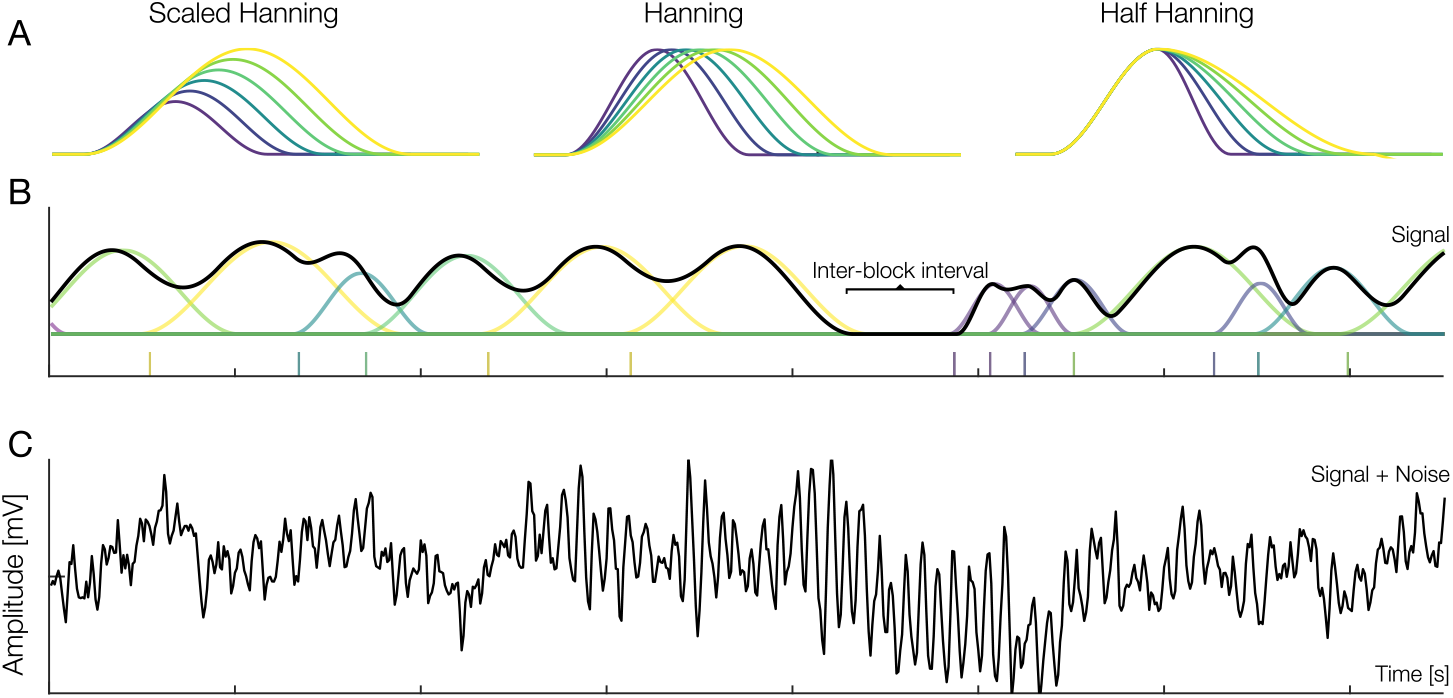
Simulation methods; (A) Three different “proto-ERPs” (i.e. ground truths). In a single simulation instance only one of the three “proto-ERPs” was used as the ground truth. The color indicates the effect of duration on the ERP. (B) A single continuous simulation with “scaled hanning” (A, left panel) as ground truth with the underlying single overlapping ERPs. Vertical bars underneath the signal indicate event onsets. Notice the perfect correlation between event distance and duration, i.e. the longer the distance between two events, the more scaled is the hanning shape; (C) the same continuous simulation with noise added.

Each simulation varied depending on multiple parameters: shape (half-hanning, hanning, scaled hanning), duration effect (yes/no), and temporal overlap (yes/no). In the case of temporal overlap, we sampled inter-event distances either from a uniform (from=0, to=3.5) or a half-normal (mu=0, std=1, truncated at 0) distribution. The same sampled inter-event distances were used as factor for the duration effect (if present), making duration effect and temporal overlap perfectly collinear. We chose these distributions as the former mimics experimentally set stimulus durations, while the latter mimics fixation durations from a free viewing task. For each combination, we generated 500 individual “proto-ERPs”. These ERPs were then added together based on the simulated onsets to get one continuous EEG signal and the accompanying event latencies (see Figure 3.B for the individual ERPs of one simulation with overlap and the corresponding continuous signal).

Additionally, ERPs were grouped together into blocks, each block containing activity from 25 ERPs, with inter-block-intervals containing no ERP activity (Figure 3.B), similar to an fMRI block design, but with the stark distinction that we analyze it as an event-based design due to the higher temporal resolution of EEG. We decided on simulating these blocks only after finding a discrepancy in results between initial simulations, which didn’t incorporate inter-block intervals and our real co-registered eye-tracking/EEG data. As such, the inter-block-intervals in our simulations mimic inter-stimulus intervals between stimuli of a free-viewing task. For a more thorough discussion on this point, why this is important, and its effects on the results, please see section Section 2.2.6.

#### 2.1.1. Noise

Subsequently, for a single simulation we produced real EEG noise by randomly selecting a minimally pre-processed closed eyes resting-state recording from one of 64 channels of one of 11 participants of a previously recorded study (Skukies, 2020, the data can be obtained from Bajwa et al., 2024; for specifics on the preprocessing see appendix Section A.1; see Figure 3.C for simulated data with added noise). We choose to use eyes-closed instead of eyes-open to not introduce additional eye movement and/ or blink artifacts into the data, which would have required further preprocessing, for example with ICA. Additionally, since the recordings tended to be shorter than the simulation, recordings were artificially prolonged by repeating the chosen recording until it matched the length of the simulation. After adding the noise, the data (i.e simulated data + noise) were FIR filtered at a −6db cut-off frequency of 0.5Hz. We filtered the data because in the overlap condition we saw a steady-state DC offset, a result from our positive-only proto-ERPs. While our deconvolution models successfully removed these offsets, the traditional averaging analysis does not. But such a large DC offset strongly influences the MSE compared to the “true” proto-ERP shape. Filtering removes this offset, but is conservative as it biases in favour of the classical averaging analysis. All simulations were repeated without noise as well, however since the main results did not change, we do not further discuss them here (see appendix Section A.2 for results without noise). Based on experience and the fact that we only minimally process the noise recordings, we judge the noise level to be realistic, and rather on a more noisy level (i.e. low SNR). Additionally, it should be remarked that our simulations pertain to the results of a single subject analysis, but our free-viewing example results (see Section 3) are based on a group of subjects, greatly increasing the SNR by aggregating results over subjects.

#### 2.1.2. Analysis

We analyzed the generated continuous time series using rERP linear overlap correction based on multiple regression. We varied three analysis parameters: overlap correction (yes/no), modeling duration (simple averaging vs. to model duration), and type of duration-modeling (four different approaches to model duration, see below), effectively forming a 5×2 design of analyses. However, the classical averaging was only calculated in order to compare duration modeling to the current standard practice of not modeling duration at all, as such we show all results relative to the classical averaged ERP (Wilkinson Formula: *y* ~ 1), which ultimately left us with a 4×2 design (overlap correction x type-of-model).

For the first approach, we introduce a linear duration effect to model duration (Wilkinson Formula: *y* ~ 1 + *dur*, termed duration-as-linear).

For the second approach, we sorted the event durations into 10 equally sized bins, and estimated a separate ERP for each of the bins, (Wilkinson Formula: *y* ~ 1 + *bin*(*dur*), termed duration-as-categorical). This was done as it, at first sight, is a simple solution to modeling varying event durations, even though it is heavily criticized by in the statistics community as “dichotomania” (Greenland, 2017), and especially in pharmaceutics and the medical field (Giannoni et al., 2014; Senn, 2005). In our case, we introduce arbitrary bins, and thus unrealistic discontinuities at the event duration bin edges. Yet, this practice has previously been presented as a favorable analysis over classical ERP averaging, and leading to an increase in power (Poli et al., 2010).

Third and fourth, we use the generalized additive modeling framework (GAM) and use a regression B-spline (please see Ehinger & Dimigen (2019) for a more practical, and Smith & Kutas (2015a) for a more theoretical explanation on the use of GAMs and B-splines within EEG) with two different levels of model flexibility (Wilkinson Formula: *y* ~ 1 + *spline*(*dur, df* = 5), and more flexibly: *y* ~ 1 + *spline*(*dur, df* = 10), termed duration-as-5spline and duration-as-10spline). There doesn’t exist a physiological explanation to choose exactly these number of splines. The number of splines solely depends on a trade-off between model flexibility and over fit, and currently there is no method implemented to estimate this automatically. Here, we chose a lower (5) and higher (10) number of splines to showcase the trade off between model flexibility and overfit, as we expected the 10 spline model to show more variance in MSE results and/or worse MSE values due to overfit.

While a few model estimations were evaluated qualitatively by plotting their time courses (Figure 4), the full set of simulations was evaluated by computing the mean squared error (MSE) between the respective model predictions and the ground truth at the 15 quantiles of the duration predictors. Subsequently, MSE values were normalized against the model with no duration effect, that is, the classical averaged ERP. This means that resulting normalized MSE values < 1 indicated that a model was performing better than classical averaging, while values > 1 indicated worse performance. In addition to the five modeling approaches, we tested the influence of actively modeling overlap using linear overlap correction using the FIR-deconvolution approach popularized through fMRI (Ehinger & Dimigen, 2019; Smith & Kutas, 2015b), against the passive method to try to resolve overlap via the duration coefficients (Nikolaev et al., 2023).

**Figure 4.**
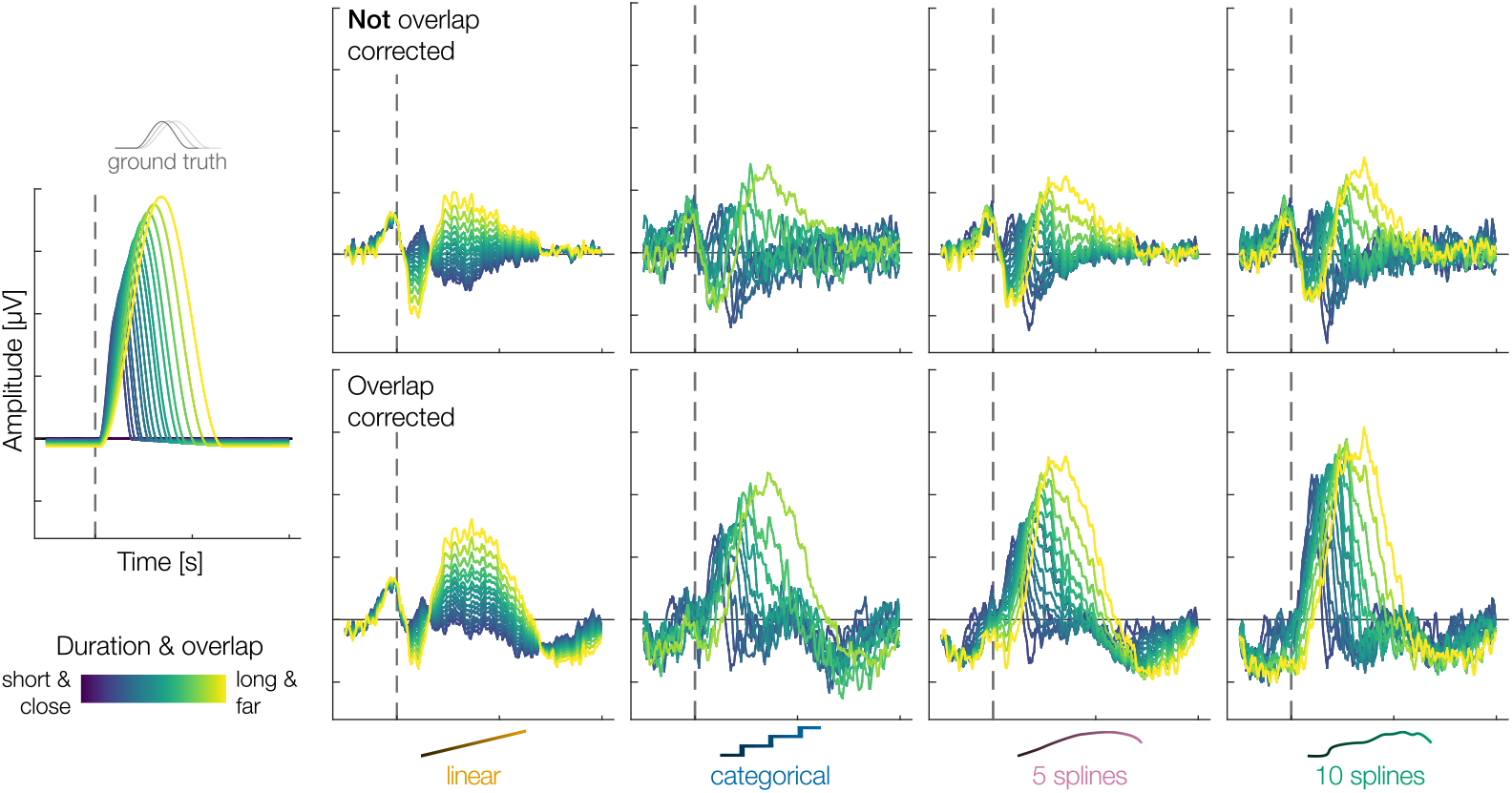
Results from a single simulation for the four tested models (including duration as: linear, categorical, 5-spline, 10-spline variable) combined with (bottom row) and without (top row) overlap correction. In this single simulation a combination of overlap correction and duration-modeling through 10-splines results in the best estimation of the ground truth(bottom row, right most panel). Left panel = ground truth. Simulation parameters: shape = scaled hanning; duration simulated = true; distribution = half-normal; noise = true; overlap = true

The above procedure, i.e. every possible parameter combination, was repeated 50 times, resulting in a total of ~15.000 simulations.

Given that statistical precision can be arbitrarily increased by increasing the number of simulations, we opted to forgo statistical hypothesis testing for the simulations entirely, other than calculating the MSE of the simulation results. We further do not report any mean MSE values, as we found them quite dependent on initial conditions and less relevant compared to the qualitative results visible. As such, we only consider conditions that are obviously different, e.g., by non-overlapping boxplots as of interest.

### 2.2. Results

We simulated ERPs with varying event durations and tried to recover the ground truth using four different modeling strategies: duration-as-linear, duration-as-categorical, duration-as-5splines, and duration-as-10splines.

In Figure 4, we show a representative analysis of one such simulation, using the scaled hanning shape, realistic noise, and overlap between subsequent events. Two results are easily visible from these plots, foreshadowing our later quantified conclusions:

1. The overall best recovery of the ground truth is achieved in the lower row, first panel from the right, the duration-as-10 splines predictor.
2. Comparing the top and bottom rows, without and with overlap correction, shows that only with overlap correction we can get close to recovering the original shape. Thus, purely modeling duration effects, cannot replace overlap correction.

However, it is important to keep in mind that this is only a single simulation. To generalize and quantify these potential effects, we calculated the mean squared error (MSE) between each analysis method’s predictions and the ground truth for 15 different event durations. All resulting MSE values are finally normalized to the results of fitting only a single ERP to all events (Wilkinson notation: *y* ~ 1), that is, modeling no duration effect at all (Figure 5).

**Figure 5.**
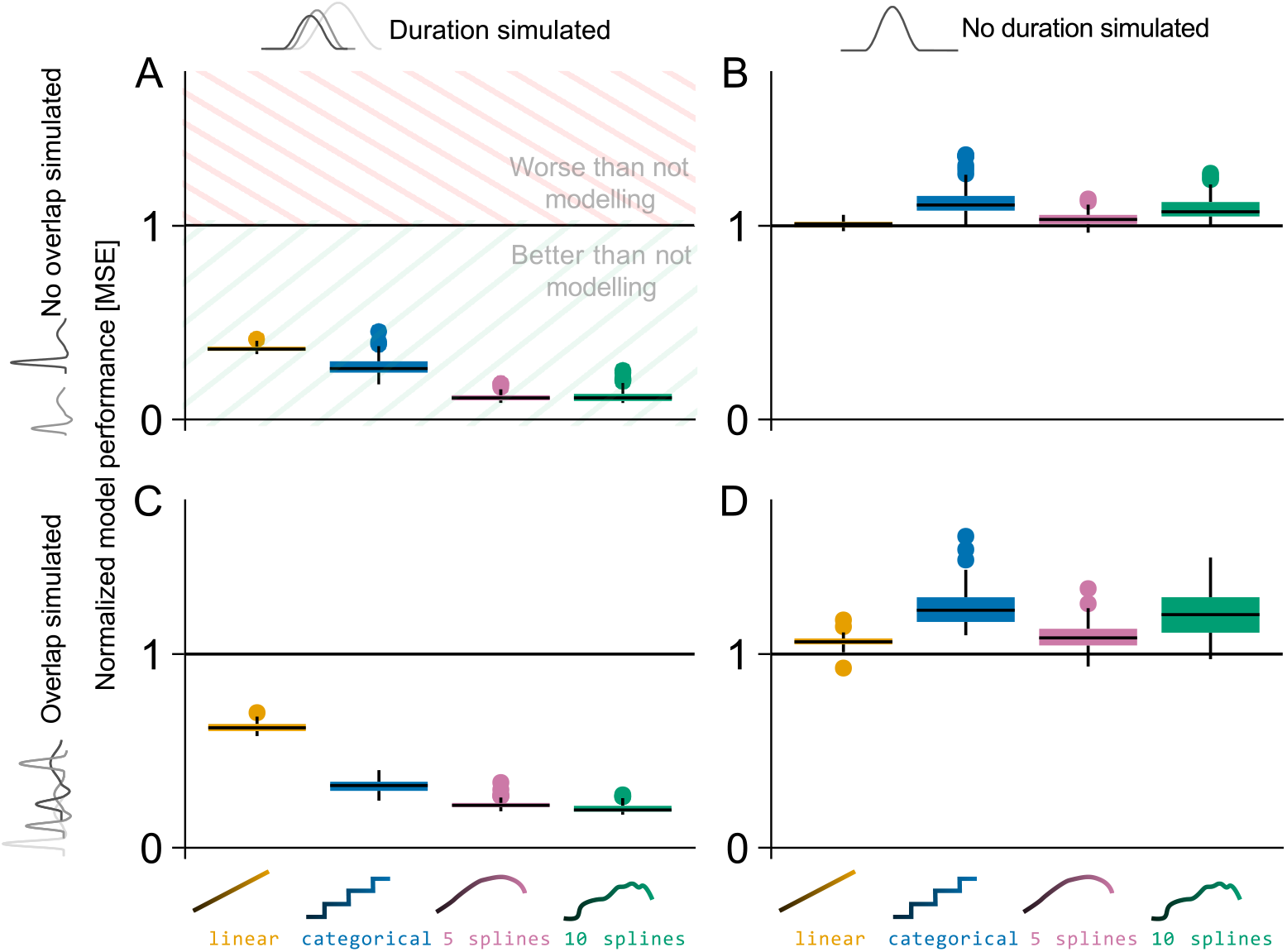
Normalized mean squared error results for the four tested models (including duration as: linear, categorical, 5-spline, 10-spline variable) in different simulation settings. Black line (y-value one) indicates results from classical averaging; MSE of zero indicates perfect estimation of the ground truth. (A) Results when a duration effect, but no overlap is simulated. The spline strategies outperform the other strategies. (B) Results when no duration effect and no overlap were simulated, but duration effects were still estimated. Little overfit is visible here. (C) Results when duration effects were simulated, signals overlap, and overlap correction was used for modeling. No interaction between duration modeling and overlap correction was observed on the MSE performance. (D) Results when a duration effect was not simulated, and signals overlap, and results are overlap-corrected.

In the following, we will present the results of the four different simulation settings, modifying both, whether we simulate overlap and duration effects, but also whether we use overlap correction in the analyses. To keep things (relatively) simple, we will first present these results by looking at simulations from only one shape (scaled hanning, see Figure 3.A) and one duration sample distribution (half normal). However, the general conclusions hold true regardless of shape and sampling distribution, as discussed later.

#### 2.2.1. The Best Suited Way to Model Duration Effects

First, we simulated duration effects without any overlap and analyzed them using several duration-modeling approaches to investigate their performance compared to not modeling the duration at all (equivalent to a single averaged ERP). Figure 5.A shows the smallest relative MSE for the spline basis with 5 or 10 splines. This indicates, foremost, that modeling duration effects is superior to not modeling it, but also shows that using splines to model duration effects is superior to using a linear or categorical duration predictors.

To check for potential overfit, we repeated the simulation without simulating any duration effect, where the normalizing (classical averaging) model should perform best. Indeed, this is visible in Figure 5.B, and, as expected with a more flexible model, we also see a small indication of overfit in the linear and non-linear model fits, but importantly with a negligible size.

Taken together, the drawback of slight overfit in cases with no duration effect is in no relation compared to the benefit of being able to capture the variability a duration effect can introduce.

#### 2.2.2. Can we Model Duration Effects and Overlap Simultaneously?

Next, we introduced overlap between events, mimicking the process of a response ERP following a stimulus ERP, or more appropriate to our simulation setup, subsequent overlapping FRPs in a co-registered EEG/ET experiment. Given that shorter event durations always co-occur with more overlap, and the event distance describes both event durations and overlap, it is not apriori clear whether both can be disentangled simultaneously.

In the case of simulating duration and overlap, and analyzing with overlap correction (Figure 5.C), we again find the result that duration-as-splines performs best. Indeed, even in our high-noise regime, the more flexible spline conditions outperform the others, without showing higher variance. This is direct evidence that overlap correction and duration-effect modeling can be used concurrently, even though the amount of overlap and the duration effect is strongly correlated by design and in nature.

Additionally, even in cases where ERPs overlap and we would wrongly assume an effect of duration (i.e., modeling duration, while no duration effect is present in the data) model performance only slightly decreases when we correct for the overlap (Figure 5.D) compared to cases where no overlap is present (Figure 5.B). This indicates that, while researchers should still carefully consider if an effect of duration is present in their data, the potential drawbacks of not modeling an existing duration effect are far greater than the opposite.

#### 2.2.3. Can we model Overlap via Duration Effects?

As a last check, we wanted to test a proposition by Nikolaev et al. (2016) and Van Humbeeck et al. (2018) in that overlap effects can be accounted for using only non-linear splines for the duration predictor. Their idea is that, because the overlap is a direct result of the duration of a trial, by keeping the effect of duration constant in the presentation of the results, one effectively adjusts for the overlap effect as well, without ever explicitly modelling overlap. However, the authors never tested this assumption through simulations. As visible in Figure 4 (compare overlap-corrected and not overlap-corrected), this seems to be insufficient. We tested whether one can use durations to model the overlap more systematically (depicted in the appendix Section A.3), and in fact, in our simulations, a model without any overlap correction or duration modeling (i.e. taking the average; Wilcoxon formula: *y* ~ 1) even outperforms modeling overlap via duration. This shows us that we indeed have to, and can, rely on other techniques like linear overlap correction, as proposed in this study.

#### 2.2.4. Interaction of Filter and Overlap Correction

We noticed an interaction between our use of a high-pass filter and the linear overlap correction, resulting in a DC-offset in the overlap-corrected estimates. This is visible in Figure 4 (the offset in the baseline period in the overlap-corrected results), and becomes more apparent when we compare non-standardized MSE values with and without overlap correction for the intercept-only model (see appendix Section A.3). Given that we know overlap correction works in the overlap simulated, but no duration effect simulated case, we would have expected overlap-corrected models to perform much better (expressed by lower MSE values) than not overlap-corrected models. And in fact, once we corrected for the baseline offset, MSE values for overlap-corrected results showed exactly this improvement (notice how MSE values improve in the first two figures in appendix Section A.4 when compared with Figure 5.C and Figure 6, and compare MSE values for the intercept-only models in appendix Section A.3 and the third figure of appendix Section A.4). However, not filtering would have resulted in strong baseline problems in the non-overlap corrected mass univariate case. We decided to present the results in the main part of our paper without any baseline correction, as this is the more conservative case, since a baseline correction would further bias results against the mass-univariate models. Moreover, it is not entirely clear whether or how baseline correction should be done in the first place, especially given that in overlap paradigms there often is no “activity-free” baseline period (for the latter see Nikolaev et al., 2016, and Alday, 2019).

**Figure 6.**
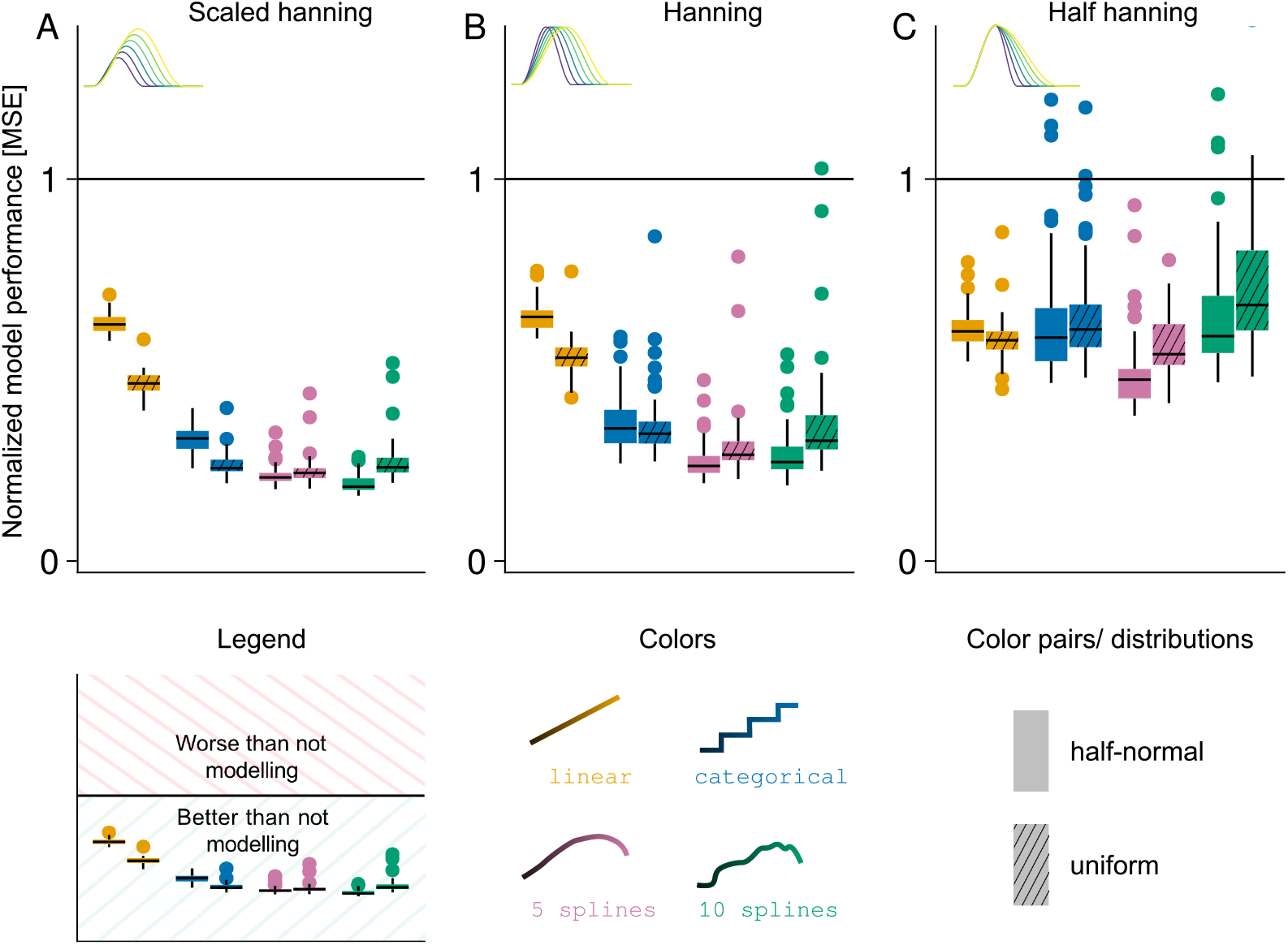
Comparison of normalized mean squared error results for the four tested models (including duration as: linear, categorical, 5-spline, 10-spline variable) between shapes (A-C) and duration distributions (color pairs). The Black line (y-value one) indicates results from classical averaging; A MSE of zero indicates perfect estimation of the ground truth. Parameter settings for all panels: duration affects shape; overlap simulated; overlap-corrected/ modeled.

At this point, we cannot say for certain where this interaction stems from, however, conducting a thorough investigation of this phenomenon is out of the scope of this paper.

#### 2.2.5. Influence of Shape and Overlap Distribution

There are two more parameters of our simulations that we have not yet presented: the overall shape of the ERP (not only the duration part), and the distribution of durations (and with that, overlap). The overall main conclusions hold true regardless of the set parameters.

Regardless of the shape of the simulated proto-ERP and overlap distribution, spline-modeling of duration consistently gives us the best results (Figure 6.A-C). However, MSE values and variance increase for the hanning shape (relative to the scaled hanning), and even further for the half hanning shape. Importantly, for the latter, variance in MSE values increased in a way, that for some simulation instances (i.e., for specific seeds) it would have been better to not model duration at all. We think that this increase in variance and value of MSE for the half hanning shape might be due to a saturation effect; if there is less variance in the duration effect to be explained in the first place, then a model without duration effect might perform similarly to a model explicitly modeling it. Lastly, we do not observe a consistent pattern of the influence of the distribution of overlap, but we cannot exclude effects in more extreme cases, due to for example highly narrow banded or bivariate distributions (Figure 6 color pairs).

#### 2.2.6. The Necessity of Block Experimental Structures

In our initial simulation of duration combined with overlap correction, we noticed a strong structured noise pattern. This pattern persisted even when no signal was simulated (Figure 7.B), and it completely disappeared when event timings from real datasets were used. We systematically tested different potential sources and found that the block structure of most experiments, that is, having a break after a set of events (Figure 7.A lower) resolves this issue. Note that this is not specific to one modeling strategy, but rather the artifact (Figure 7.B) can appear even with a linear effect. We quantitatively tested this finding and show that artificially introducing a break after every 25 events (which results in only 20 breaks in our simulations) completely removes this artefact (Figure 7.B vs. Figure 7.C, Figure 7.D).

**Figure 7.**
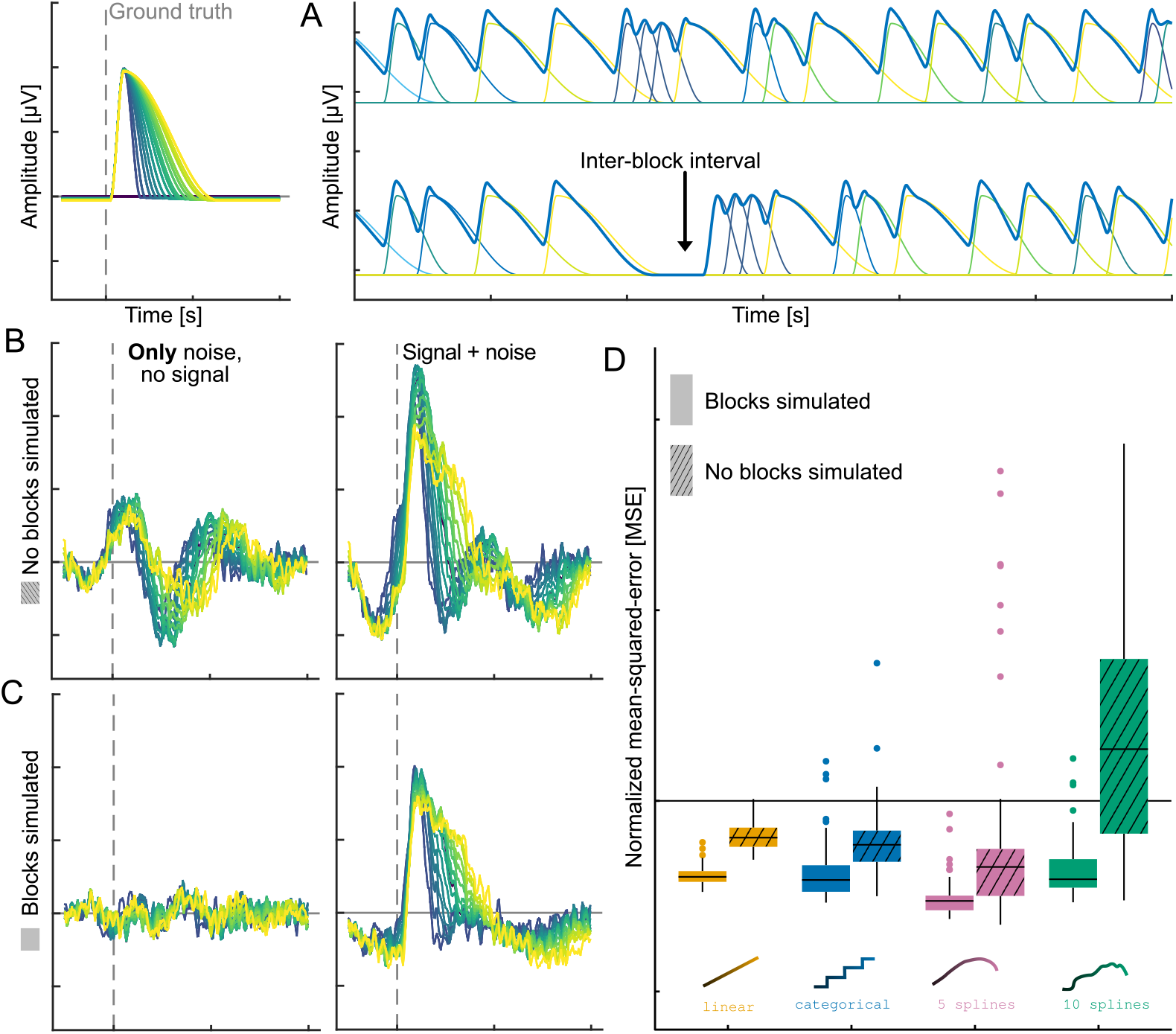
Comparison of simulations with and without simulated block structure. Up left: Ground truth shape “half hanning” used in the depicted simulation; (A) Continuous simulated EEG and its underlying responses without noise, once without (up), and once with (down) added blocks; (B) Estimates from one simulation without added blocks, once without any signal in the data (left, i.e. the data contained only noise) and once with added signals (right, i.e. the data contained noise + signal); (C) Estimates from the same simulation, but with added blocks every 25 events, once without any signal in the data (left, i.e. the data contained only noise) and once with added signals (right, i.e. the data contained noise + signal); (D) Comparison of MSE results between simulations with blocks added (full colors) and simulations without any blocks added (cross-hatched colors) for all tested models in conjunction with overlap correction.

We assume that including the duration of events both as a predictor and explicitly in the FIR time-expanded designmatrix (as inter-event distance) may lead to a specific type of collinearity. This collinearity significantly compromises the reliability and stability of the model estimates, and due to the duration effect, results in smooth wave-like patterns (Figure 7.B). When introducing (seemingly) overlap-free events at the beginning and end of a block, this collinearity between the duration and the structure of the designmatrix is broken, and a unique solution can be estimated.

## 3. Real Data Example 1: EEG + ET

We move our eyes approximately 120-240 times each minute. Besides being an interesting behavior in itself, eye movements offer a unique and data-rich segmentation event for EEG. Consequently, more and more researchers use EEG and eyetracking to better understand human cognition (Dimigen et al., 2014; Dimigen & Ehinger, 2021; Nikolaev et al., 2016). Here, EEG and eye movements are co-registered, such that single fixations can be used as event markers in the EEG. These fixations can then be assigned to different conditions based on the fixated stimulus, and modeled accordingly to achieve a fixation-related potential.

To illustrate the interplay of duration and overlap modeling, we use a public co-registered unrestricted viewing dataset by Gert et al. (2022). During the study by Gert et al. (2022) participants were presented with faces in one (passive) condition, where participants were told to fixate a fixation cross. Natural scenes were presented in a second (active) condition, where participants were allowed to freely explore the presented stimulus through eye movements. We analysed only the second condition containing natural scenes. Each participant was presented with 171 natural scene images, presented at a size of 30.5 × 17.2 degrees angle. Each trial consisted of a fixation circle (presented for 1.8-2.2 sec.), followed by a natural scene (presented for 6 sec.), followed by a blank screen (presented for 1.6-2.0 sec.). For a visual presentation of the experiment please see Figure 1 of Gert et al. (2022).

### 3.1. Methods

The data by Gert et al. (2022) was already preprocessed as follows: First, the eyetracking and EEG data were integrated and synchronized using the EYE-EEG toolbox (http://www2.hu-berlin.de/eyetracking-eeg) (Dimigen et al., 2014). Next, the data was down-sampled from 1024 Hz to 512 Hz and high-pass filtered at 1Hz (−6 db cutoff at.5 Hz). Muscle artifacts and noisy channels were manually inspected and marked or removed respectively. Subsequently, ICA was computed, and artefactual components were subtracted. Lastly, the data was re-referenced to an average reference and removed channels were interpolated through spherical interpolation. For more specifics on the preprocessing, please see Gert et al. (2022).

To showcase our approach, we then calculated fixation-related potentials (FRP) using custom scripts and the Unfold.jl toolbox (Ehinger & Alday, 2024). The data was modeled using a linear model with fixations following the Wilkinson notation:

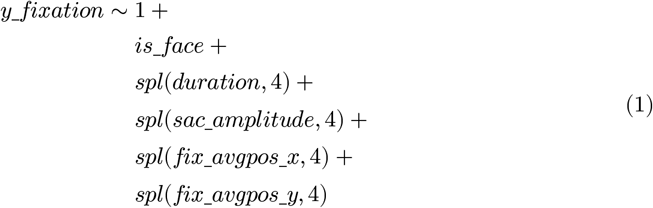

Where *y_fixation* is the recorded activity related to a fixation, *is_face* represents whether in that fixation a face was fixated (yes/no), *duration* the fixation duration; *sac_amplitude* the saccade amplitude; *fix_avgpos_x & fix_avgpos_y* the average x- and y-position of the current fixation. Apart from fixation duration, we incorporated the other covariates in the analysis as they are known to have an influence on the FRP (Gert et al., 2022; Nikolaev et al., 2023), and we wanted to adjust for these factors when investigating fixation duration. Additionally, the linear model lets us incorporate a different basis function to model stimulus onsets (i.e. the appearance of a picture on screen; for an explanation on how different basis functions for different events are combined in one linear model please see Ehinger & Dimigen, 2019) using a

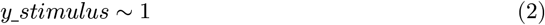

formula, as the onset of a stimulus can have huge overlapping effects on subsequent fixations (Coco et al., 2020; Dimigen & Ehinger, 2021; Gert et al., 2022). Incorporating the stimulus onset in our model in this way lets us account for the overlap the stimulus response has with any subsequent fixation.

As in our simulations, the model was calculated once with, and once without overlap correction. Because we were only interested in the effect of duration, we then calculated marginalized effects of duration on the ERP, while holding all other covariates constant at their respective means.

Finally, we statistically tested for the duration effect with and without overlap, and for the difference of the duration effect between with and without modeling. As proposed in Pernet et al. (2011), we use a Hotelling T^2^ test (implemented in HypothesisTests.jl v0.11.0), but on the spline coefficients to test them against the null hypothesis of all coefficients being 0 across subjects. We subsequently calculate a Benjamini-Yekutieli, via MultipleTesting.jl (Benjamini & Yekutieli, 2001; Gehring et al., 2023)), false discovery rate correction over all channels and time points to address the multiple comparison problem. To assess the difference, we examined the disparity in the maximal marginal effects (calculation described below) between without and with overlap correction using an FDR corrected one sample t-test. Note that the FDR correction only controls weak family wise error and direct interpretation of time and location can be problematic (Rousselet, 2025; Winkler et al., 2024).

For visualization, we estimate marginal effects over a range of durations (−0.1 to 1s). To visualize our results in a topoplot series (Mikheev et al., 2024a), we break down the multi-parameter spline estimate to a single value per channel and time-point, we calculate the maximal marginal effect over the range of durations calculated before. This quantity is always positive (the maximum needs to be >0). We therefore decided to test the coefficients directly (as described above), rather than the marginalized effects we visualize. This has the unfortunate effect that in some cases, what is tested is not the same as what is visualized, the reason being the lack of visualization tools for non-linear effects. Only for the difference do visualization and test directly correspond to each other. Visualizations were done in Julia (Bezanson et al., 2017) using the UnfoldMakie.jl (Mikheev et al., 2024b) and Makie.jl (Danisch & Krumbiegel, 2021) libraries.

### 3.2. Results

As indicated in Figure 8, without overlap correction we find three clusters of fixation duration effects along the midline: over frontal, occipital and centro-parietal electrodes.

**Figure 8.**
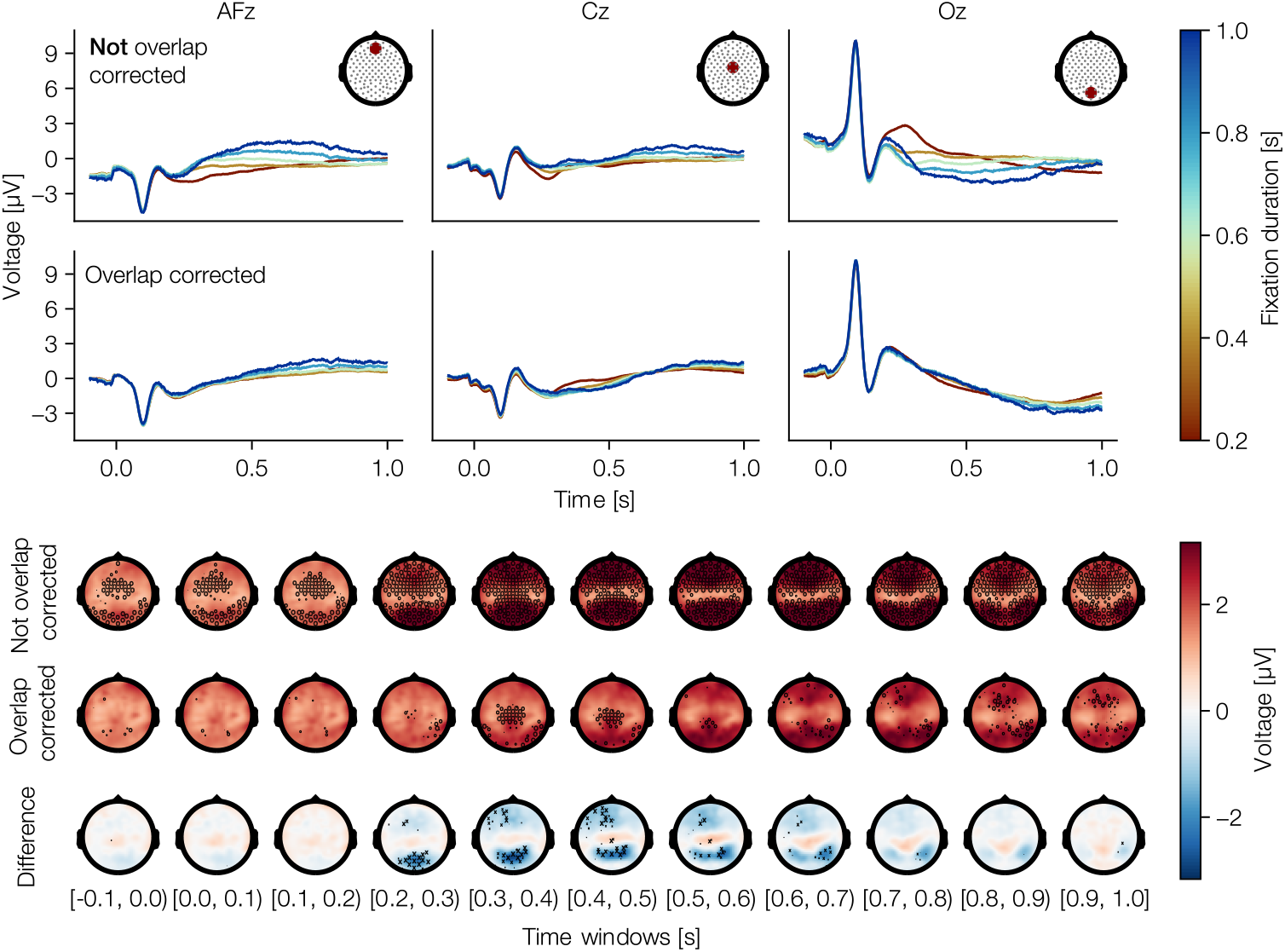
Results from a free viewing experiment. (top) Regression FRPs with marginal effects of fixation duration from channels AFz [left], Cz [middle], and Oz [right], once without [up] and once with [bottom] overlap correction. (bottom) Topoplots for results without overlap correction [up], with overlap correction [middle], and the difference between with and without overlap correction [bottom]; Displayed values are the maximal difference between two evaluated marginal effects (i.e. the maximal difference between any line in top), As a statistical test, we used a Hotellings-t^2^ test on all coefficients of the non-linear spline coefficients, not only the maximal difference, which was used only for the display. The significance of the difference was tested with mass-univariate one-sample t-tests, and corrected for multiple comparisons using FDR. We plot a marker, only if the minimal corrected p-value in a time window is below 0.05 and weight its size logarithmically according to the p-value (that is, larger markers indicate smaller p-values).

However, when modeling overlap in addition to duration, we found duration effects to become smaller and even vanish. This indicates, as proposed by Nikolaev et al. (2016), that a duration predictor tries to compensate for the overlap (but ultimately fails to do so; see simulation results Section 2.2.2). Yet, an effect over central electrodes at around 0.450 s. after fixation onset still persists in the results after overlap correction (Figure 8). This is direct evidence for the existence of event duration effects in EEG-eye-tracking co-registered data. Finally, it should be noted here that these results seem much less noisy than our simulation results Figure 4. The reason being that here we aggregated over a group of subjects, whereas our (single) simulation results essentially compare to a single subject analysis.

## 4. Real Data Example 2: fMRI

To showcase that our approach generalizes to modalities other than EEG, we reanalyzed a subset of the fMRI data used by Mumford et al. (2023), their Stroop task.

### 4.1. Methods

The dataset consists of 110 subjects completing a Stroop task (Stroop, 1935). During the Stroop task, participants are asked to name the font color of stimuli that are either congruent (e.g. the word green written with the font color green) or incongruent (e.g. the word green written in the font color red). Participants were presented with 96 trials and exhibited a mean reaction time of ~0.691 seconds. Further details can be obtained by contacting the authors of Mumford et al. (2023). fMRI signals were acquired using single-echo multi-band EPI with the following parameters: TR = 680ms, multiband factor = 8, echo time = 30ms, flip angle = 53 degrees, field of view = 220mm, 2.2 × 2.2 × 2.2 isotropic voxels with 64 slices. The data was already preprocessed using fMRIprep as described in Mumford et al. (2023):

Data were preprocessed in Python using fmriprep 20.2.0 (Esteban et al., 2019). First, a reference volume and its skull stripped version were generated using a custom methodology of fMRIPrep. A B0-nonuniformity map (or fieldmap) was directly measured with an MRI scheme designed with that purpose (typically, a spiral pulse sequence). The fieldmap was then co-registered to the target EPI (echo-planar imaging) reference run and converted to a displacements field map (amenable to registration tools such as ANTs) with FSL’s fugue and other SDCflows tools. Based on the estimated susceptibility distortion, a corrected EPI (echo-planar imaging) reference was calculated for a more accurate co-registration with the anatomical reference. The BOLD reference was then co-registered to the T1w reference using bbregister (FreeSurfer) which implements boundary-based registration (Greve & Fischl, 2009). Co-registration was configured with six degrees of freedom. Head-motion parameters with respect to the BOLD reference (transformation matrices, and six corresponding rotation and translation parameters) are estimated before any spatiotemporal filtering using mcflirt (FSL 5.0.9, Jenkinson et al., 2002). BOLD runs were slice-time corrected using 3dTshift from AFNI 20160207 (Cox & Hyde (1997), RRID:SCR_005927). The BOLD time-series (including slice-timing correction when applied) were resampled onto their original, native space by applying a single, composite transform to correct for head-motion and susceptibility distortions. These resampled BOLD time-series will be referred to as preprocessed BOLD in original space, or just pre-processed BOLD. The BOLD time-series were resampled into standard space, generating a preprocessed BOLD run in MNI152NLin2009cAsym space. First, a reference volume and its skull-stripped version were generated using a custom methodology of fMRIPrep. Automatic removal of motion artifacts using independent component analysis (ICA-AROMA, Pruim et al., 2015) was performed on the preprocessed BOLD on MNI space time-series after removal of non-steady state volumes and spatial smoothing with an isotropic, Gaussian kernel of 6mm FWHM (full-width half-maximum). Cor responding “non-aggresively” denoised runs were produced after such smoothing. These data were used in our time series analysis models.

— (Mumford et al., 2023)

For more details of the data collection and preprocessing pipeline, please see Mumford et al. (2023).

After fMRIPrep, we read the preprocessed functional data into Julia (Bezanson et al., 2017) using the NIfTI.jl package (v.0.6.1, 2025). Next, to simplify the analysis and increase SNR, we performed a region of interes (ROI) analysis based on the 100-parcellation of the Schaefer 2018 atlas (Schaefer et al., 2018). Based on Mumford et al. (2023), we expected duration effects to result in widespread activity across the brain. We averaged voxels inside each ROI and used a 4th-order Butterworth highpass at a −3db cutoff of 1/128s to remove slow drifts from the BOLD signal (*JuliaDSP/DSP.jl*, 2025). For the RT-covariate, we used a winsorization proceduce, effectively clipping the highest and lowest 5% RTs. RTs were then mean-centered and normalized to changes in standard deviation to align RT distributions over subjects. To fit the deconvolution model, we used an finite impulse response (FIR) basis function with 24 lags, starting from −0.68 to 14.96 seconds at a TR of 0.68s. While we found similar results using a HRF basis function with the standard SPM parameterization, we think based on prior work that a non-linear scaling of the BOLD function is not an adequate model of the underlying data (Grinband et al., 2008). To specify and estimate the models we used Unfold.jl (v0.8.4, Ehinger & Alday, 2025) and UnfoldBIDS.jl (0.3.3, Ehinger & Skukies, 2025). We expanded the FIR kernel to model the RT effect with a B-spline set of five splines, and additionally an effect of condition according to the following Wilkinson Formula:

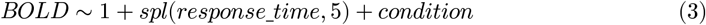

Currently, penalized splines are not defined; thus, we manually fix the number of splines to five. A higher number of splines (and thus a higher flexibility in the possible non-linearity; we tried 10 splines) resulted in strong overfit and thus much reduced significances. This was unsurprising to us, given that we only have 10 minutes of fMRI data per subject and no pooling over subject was used in the estimation process.

As in the free viewing analysis, we use Hotelling-T^2^ test to test the spline coefficients against the statistical null hypothesis of them being 0. We finally calculated a Benjamini-Yekutieli false discovery rate correction, via MultipleTesting.jl (Benjamini & Yekutieli, 2001; Gehring et al., 2023)), over the 100 ROIs and 24 timepoints.

### 4.2. Results

Mumford et al. (2023) established a reaction time effect in multiple different experimental paradigms. Here, we re-analyzed their dataset using a ROI approach, and successfully reproduced this wide-ranging reaction time effect for the Stroop task (Figure 9.A). Next, we replaced the linear term with a non-linearity via a spline-set. Again, we find a wide-spread reaction time effect, albeit, using a strict multiple-correction control, only significant in three ROIs. Comparing results from a model using a linear predictor with a model using a non-linear predictor (Figure 9.C), it is visible that the non-linear model captures a non-linear pattern in the BOLD time course, which a linear predictor cannot. Note that with our B-spline basis, it is not straightforward to test the superiority of the non-linear over the linear model, and we leave this for future studies. Further note, that we have very little data (10min) to estimate such a non-linear effect, which resulted in lower power, as indicated by larger p-values (Figure 9.B). In summary, we succeeded in our main goal to demonstrate the feasibility of combining overlap correction and duration modelling for fMRI data.

**Figure 9.**
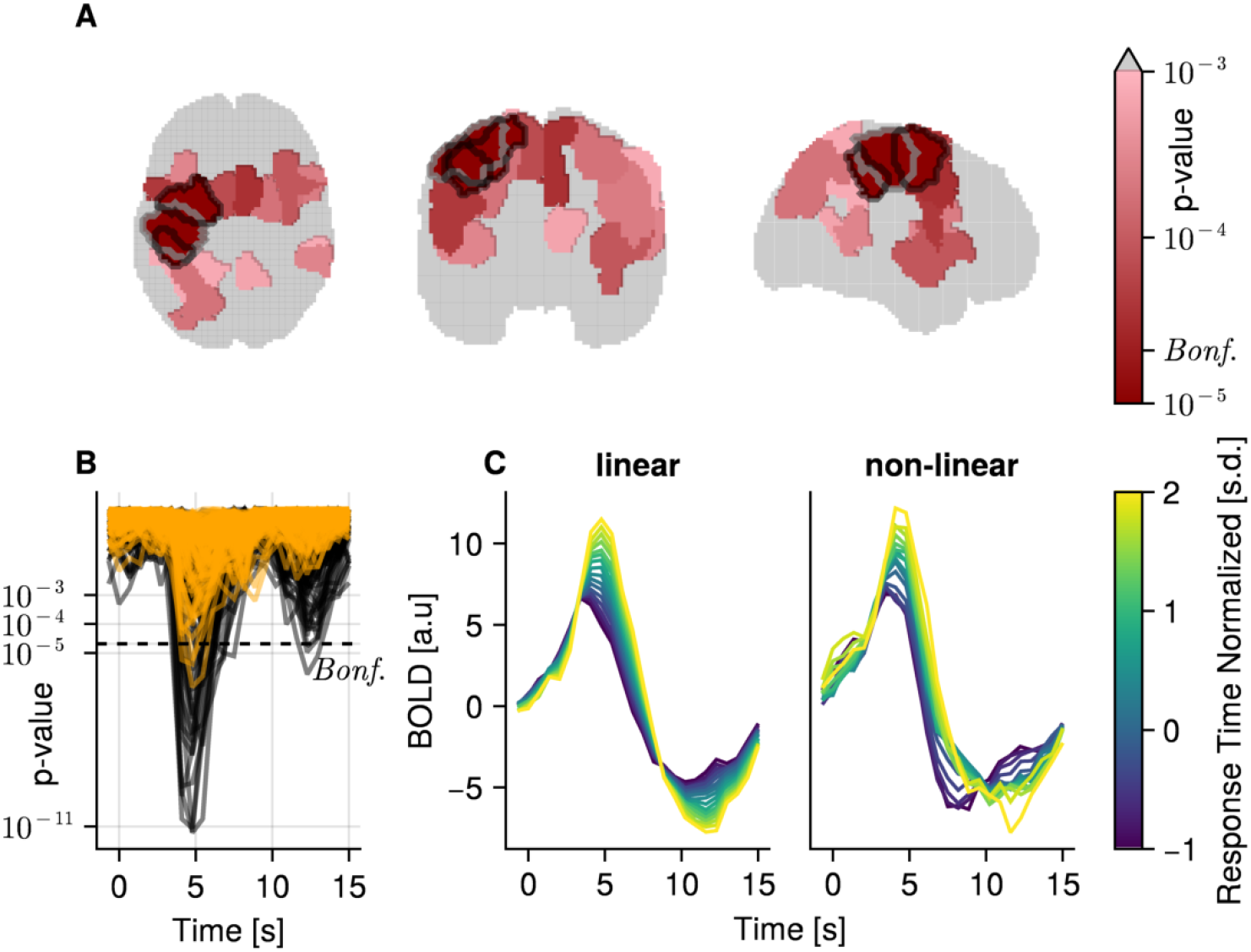
fMRI results from a stroop experiment. (A) One slice of the transverse, coronal, and sagittal planes with ROIs colored according to their minimal p-value over time, uncorrected. Grey areas indicate p-value > 10^−3^. Indicated Bonferroni threshold was calculated over time and 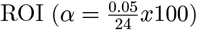. Marked regions (black outline) were significant under FDR correction over time and ROI. (B) Uncorrected p-values over time for all ROIs, with a linear model (black) and non-linear model (orange). The dashed line indicates the cut off for p-values after Bonferroni correction. (C) Marginal effects of the estimated BOLD response of one example region (gray outlined in A), for the linear (left) and non-linear (right) model.

## 5. Discussion

Event durations often differ between conditions and participants. While in some cases this can be controlled, in many cases it cannot. As soon as the event duration is under the control of the subject (e.g., differing reaction times in old vs. young, or with the recent trends towards quasi-experimental designs; Gramann et al., 2021, Welke & Vessel, 2022, Mathewson et al., 2024), many aspects of an experiment are no longer under the control of the experimenter. Here, we first show that such a discrepancy can lead to spurious effects in the estimated brain responses. We next demonstrate that when explicitly modeling such event duration effects, these spurious effects can be successfully addressed. Finally, we systematically evaluated that non-linear modeling of event durations is compatible with linear overlap correction and that implementing such analysis can give a better understanding of the true brain processes.

### 5.1. Modeling Event Durations

One of the key takeaways from our study is the potential for spurious effects in neural data analysis when variations in event durations are not considered. This is in line with research by Wang et al. (2018) which showed that neural activity can scale in proportion to task duration, and Yarkoni et al. (2009) and Mumford et al. (2023), that argued for including durations during modeling of fMRI data.

In EEG research, earlier studies proposed to circumvent this problem by binning reaction times and calculating the average ERP of each bin (Poli et al., 2010) (and is now a prominent suggested method in ERP analysis; see the chapter on event bins in Luck, 2022) or as a non-linear predictor via a smoothing function (Hassall et al., 2022; Van Humbeeck et al., 2018). Contrary to what the issue of “dichotomania” (Senn, 2005) might suggest at first sight, our simulations show that categorizing durations into bins performs better than even linear modeling of duration. This is a result of the underlying event duration effect not being linear but rather non-linear. To summarize, while binning is able to improve results over not modeling and linear modeling, it does so by introducing unrealistic discontinuities which can be remedied by using splines.

We show the critical role that duration plays in the interpretation of ERP results, and further, that it takes the flexibility of non-linear spline estimation via multiple regression to model event durations and achieve meaningful results. As such, to ensure accuracy and reliability of results, duration effects cannot be ignored, but rather must be addressed.

### 5.2. Compatibility with Linear Overlap Correction

The presence of overlap in EEG data is a common phenomenon. And although this problem has been thoroughly addressed before (Ehinger & Dimigen, 2019; Smith & Kutas, 2015a), it was not clear if modeling of event durations and overlap correction can be combined. That is, through a dependence of varying-event timings needed for linear overlap correction, which in turn are directly related to the duration effect, we were left with a potential collinearity problem.

Our simulations indicate that non-linear event duration modeling and linear overlap correction of adjacent events can be seamlessly integrated. We also showed that this comes at a minimal cost in terms of potential overfit, should there be no ground truth duration effect in the data. We therefore recommend using our proposed non-linear duration modeling in cases researchers suspect effects of duration.

### 5.3. Further Considerations

In addition to overlap and duration estimation method, we considered four further parameters: ERP shape, block-design, jittering inter-onset distances, and regularization.

- <u>ERP shape:</u> Overall, results do not change drastically based on different ERP shapes, i.e. it is still best to model duration by non-linear spline estimation in conjunction with overlap correction. However, while results from two of our shapes (scaled hanning and hanning) showed very similar results, the half hanning shape showed signs of increased overfit for 10-splines, with larger MSE compared to the best-performing 5-splines. Currently, the number of splines in overlap-corrected regression fits has to be chosen by hand, instead of an automatic penalized splines method due to computational costs. Similarly, the variability of MSE values within model categories increased, especially for the more flexible models. This is likely due to the shorter time window of the duration effect itself, and thus these flexible models are more prone to overfit on the noise in the earlier non-changing part of the shape. We cannot exclude more extreme shapes to show more extreme behavior.
- <u>Block-design:</u> As described in Section 2.2.6, a continuous stream of overlapping events without breaks leads to increased overfit under certain simulation conditions. Introducing “inter-block intervals”, where at least two adjacent events break the homogeneous overlap-duration relationship, alleviates this issue. We think that such pauses break the correlation between inter-event onsets as modeled with the FIR time expansion, and the duration predictor. Further simulations showed that this effect is not specific to splines, but also occurs with linear duration modeling. Further, this effect is irrespective of shape and occurs in pure noise data as well (see Figure 7). However, by adding only one or more (depending on SNR) of the aforementioned inter-block intervals, we could resolve this noise amplification. In real data such inter-block intervals can occur, for example in free-viewing experiments where FRPs are of interest. Here, while fixation duration and overlap are indeed perfectly correlated within the e.g. 5s of looking at a stimulus, the last fixation before removing the stimulus does have a duration, but no further (modeled) fixation overlapping. In other experiments, blocks can (and usually are) in the experiment by design.
- <u>Jittering inter-onset-distances:</u> In our free viewing data, we also noticed that the inter-event onsets and durations are not as perfectly correlated as in our simulation due to variable saccade duration. We therefore introduced a uniform jitter of +-30ms (expected saccade duration) to the inter-event-onsets in some exploratory simulations. This had only a minimal effect, if any at all, in these simulations. We did not test this systematically, as it is outside the scope of this paper.
- <u>Regularization:</u> Lastly, we also ran some preliminary tests on L2 regularization (Kristensen et al., 2017). Regularization should reduce overfit on the data at the expense of introducing a bias, in the case of the L2 norm, towards coefficients with smaller amplitude. Typically, the amount of regularization is chosen by cross-validation, however, in our simulations, we often observed that this resulted in regularization to (near) zero. Currently, in Unfold only a “naive” cross-validation within the cvGLMNET solver (Qian et al., 2013) is available. This method is insufficient because cross-validation splits are automatically set by a blackbox-function, and thus can be within-blocks, potentially leading to data leakage between test and validation set. We are not aware of a sufficient implementation alleviating this problem. Nevertheless, we identified a second, more severe limitation than data leakage: There is only a single regularization parameter for the whole model, thus with overlap correction, we regularize all time points relative to an event onset equally. This means that baseline periods, where the true regularization coefficient should be very high, as there is no reliable time-locked activity, wash over to periods of activity, greatly reducing their amplitude, often to zero. Setting the regularization parameter by hand, as expected, leads to a strong reduction of this noise artifact, but at the cost of bias. Only more nuanced applications of regularization can fully address this issue, and more work is needed.

### 5.4. Limitations

#### 5.4.1. Limitations of our Simulations

In this study, we focused on three rather simple shapes for simplicity’s sake. These three shapes cover a range of possible changes to the shape in that they stretch in time, both in time and amplitude, or only partly, which are simplifications of real examples (Hassall et al., 2022; Wang et al., 2018). However, although earlier simulations with more complicated shapes that included multiple positive and negative peaks showed no substantial differences in results, we cannot be certain how our results generalize to all possible shapes.

Furthermore, our study is in several cases limited to relative comparisons between analysis methods, where the spline-modeling performed best. This does not mean that it ultimately recovers the underlying shape perfectly. Indeed, inspecting Figure 4, one can clearly see that the pattern is not an exact recovery, but confounded by noise - still, it is performing much better than ignoring the issue and only using a single ERP. Similarly, we use only a single measure (MSE) to evaluate and delineate the different analysis methods, other choices of measures are possible (e.g. correlation, absolute error, removal of baseline period). We do not expect qualitatively different results.

Next, we did not investigate if there are interactions between a condition effect and the duration effect. That is, throughout the paper, we assume that the duration effect is the same in all conditions. If we allow for such an interaction, then it is unclear to us, if these per-condition-duration-effects could be used to explain the main effects of conditions as well.

Additionally, our simulation study covers a large area of simulation parameters, but cannot cover all possible cases. Indeed, some interesting variations could be to introduce continuous and categorical condition differences, incorporate multiple event types, use more flexible non-linear analysis methods (e.g. DeepRecurrentEncoder, Chehab et al., 2022) or try to simulate empirically observed duration effects more realistically.

Lastly, we had a coarse look at different noise levels (unreported), and two noise types (white and real noise). Both factors did not influence our conclusions or the relative order of the methods.

#### 5.4.2. Limitations of Results

Another clear limitation of our study is, that we only look at model performance of what essentially constitutes results from a single subject. In this paper, we do not address how the performance will improve if estimates over multiple subjects are aggregated. In the applied example, we average results over multiple subjects, but here we want to highlight a potential caveat relating to spline modeling in general: Experimenters need to decide whether to normalize durations per subject, effectively modeling quantiles over subjects, or assume that ERPs relate better to absolute durations. Because knot placement and coverage of splines depend on the distribution of each single subject, not all subjects have the same smoothing factor. Interested readers are pointed to (Wood, 2017) for a discussion of these issues.

A more general limitation is, that we cannot know how such duration effects are implemented in the brain. This could potentially lead to an issue of causality, as the duration covariate is used to explain points in time at which the system has not yet actually specified the event duration. This limitation is not specific to this study, but in general whenever behavioral responses are used to predict neural data.

Lastly, the performance of our method compared to the approach by Hassall et al. (2022) remains to be shown. They assume a single response is stretched in time, setting a strong prior on the possible relation. Such a relation would be fulfilled in our simulation for the stretched hanning window, but not in the other shapes.

### 5.5. Application to Real Data

In our paper, we apply our approach to two datasets from two different modalities: EEG and fMRI. In both datasets we demonstrate a significant effect of duration (fixation duration for EEG, reaction time for fMRI).

#### 5.5.1. EEG

In direct comparison to our simulation results, we first applied our approach on FRPs in an unrestricted viewing dataset. Given that fixations differ in durations, with an average of 230ms, but up to a second, we expected that the evoked potential is modulated by duration as well. And indeed, without overlap correction, we clearly found strong non-linear duration effects. However, most of the effect was better explained by overlap rather than duration effects. Subsequent analysis still showed a significant duration effect, but it was comparatively small in absolute effect size. This was surprising as one earlier study (Ehinger et al., 2018; Nikolaev et al., 2023) found a larger effect size of duration for a similar analysis. To better characterize and understand this effect, more datasets should be analyzed; nevertheless, we can conclude that we successfully disentangled overlap from fixation duration effects.

#### 5.5.2. fMRI

In Section 4 we directly show that our approach is applicable to functional magnetic resonance imaging (fMRI). fMRI has the advantage that the underlying measure generator is relatively well understood (as the hemodynamic response function; HRF). As such, the problem of overlapping signals has been thoroughly addressed (Penny et al., 2011). Yet, while some studies have shown the importance of linearly modeling reaction times in fMRI studies (Mumford et al., 2023; Yarkoni et al., 2009), this is not common practice yet. Adding to this, we show in Section 4 that one can (and should) model reaction time via non-linear spline regression instead of linear estimation. Further, both studies only focus on reaction time, whereas we argue to generalize such modeling to other event durations, such as stimulus durations or fixation durations.

One possible different approach to deal with the problem of varying durations was proposed by Grinband et al. (2008). In their study, the authors come up with two different models incorporating duration to estimate the BOLD signal: One model uses “variable duration epochs” (duration modulated boxcars) to convolve the HRF; the other uses the more common constant impulse response function (IRF) with an additional regressor linearly modulated by duration (Figure 9.C, black lines). The authors found that both their approaches increased statistical power, consistency, and interpretability of results. However, these models are only able to either capture the time-aspect (variable epoch) or the amplitude-aspect (constant IRF with regressor) of the duration effect. Our approach improves upon their models in two ways: First, our approach is able to capture duration effects in both time and amplitude. And second, our model is able to incorporate non-linear effects, a refinement that was already proposed by Grinband et al. (2008).

### 5.6. Generalization to other Modalities

While in this paper, we focus on applying our approach to M/EEG data and further showcase it on fMRI data, the technique can be extended to other neuroimaging modalities. In fact, it is plausible that almost all brain time series analyses could benefit from considering event duration as a potential confounder. Here we give two examples.

First is the investigation of local field potentials (LFP). This should come as no surprise, as LFP are inherently very close in analysis to EEG. That is, LFPs consist of fast-paced electrical fields which might overlap in time and are likely subject to change depending on the length of an event. In fact, at least one study in non-human primates already highlighted the relationship between task timing and LFPs (Kilavik et al., 2010). However, their task involved the primates to execute a limb movement of one of two interval lengths, ultimately resulting in two conditions (i.e. long vs. short interval) with multiple trials over which were then aggregated, instead of considering duration as a continuous predictor. Additionally, overlap was not considered.

Secondly, pupil dilation measurements could benefit from our analysis approach as well. Typically, pupil dilation is taken as an indirect measurement of arousal states, cognitive load, and has been linked to changes in the noradrenergic system (Larsen & Waters, 2018). Whereas pupil dilation has often been related to reaction time (Hershman & Henik, 2019; Isabella et al., 2019; Strauch et al., 2020), to our knowledge no study so far has modeled reaction time explicitly. At least one theory-driven model has been proposed which offers explanations for a wider range of parameters, including trial duration (Burlingham et al., 2022). However, again overlapping signals are dismissed in this model, and as such our more general approach is to be preferred.

## 6. Summary

In the present study, we show through extensive simulations that event durations can and should be modeled in conjunction with overlap correction. Through combining linear overlap correction with non-linear event duration modeling, researchers gain a more powerful tool set to better understand human cognition. Overlap correction disentangles the true underlying signals of the ERPs, while event duration modeling can lead to a more nuanced understanding of the neural responses.

By applying this analysis to real data, we underscore the importance of modeling event durations both, in cases where they are of interest themselves, as well as in situations where durations may have an interfering influence on the results.

As such, we advise researchers who study brain related time series data to take special care to consider if and how overlap and event durations affect their data.

## Data and Code Availability

All code is publicly available at https://github.com/s-ccs/unfold_duration. The FRP data can be obtained from Gert et al. (2022). Raw fMRI data can be obtained from https://openneuro.org/datasets/ds004636/, for the preprocessed data please refer to the corresponding author of Mumford et al. (2023).

## Author Contributions

- <u>René Skukies:</u> Conceptualization; Methodology; Software; Formal analysis (simulations, fMRI analysis); Data Curation; Writing - Original Draft; Visualization
- <u>Judith Schepers:</u> Formal analysis (FRP analysis); Writing - Review & Editing
- <u>Benedikt Ehinger:</u> Conceptualization; Methodology; Software; Resources; Supervision; Formal analysis (simulations, FRP analysis, fMRI analysis); Writing - Review & Editing; Visualization; Funding acquisition

## Acknowledgements

We want to specifically thank Jeanette Mumford for sharing her pre-processed fMRI dataset with us.

## Bibliography

Alday, P. M. (2019). How Much Baseline Correction Do We Need in ERP Research? Extended GLM Model Can Replace Baseline Correction While Lifting Its Limits. Psychophysiology, 56(12), e13451. 10.1111/psyp.13451

Bajwa, I. J., Nilsen, A. S., Skukies, R., Aamodt, A., Ernst, G., Storm, J. F., & Juel, B. E. (2024). “A repeated awakening study exploring the capacity of complexity measures to capture dreaming during propofol sedation” [Computer software]. OpenNeuro. 10.18112/openneuro.ds005620.v1.0.0

Benjamini, Y., & Yekutieli, D. (2001). The Control of the False Discovery Rate in Multiple Testing under Dependency. The Annals of Statistics, 29(4), 1165–1188. 10.1214/aos/1013699998

Bezanson, J., Edelman, A., Karpinski, S., & Shah, V. B. (2017). Julia: A fresh approach to numerical computing. SIAM Review, 59(1), 65–98. 10.1137/141000671

Brands, A. M., Devore, S., Devinsky, O., Doyle, W., Flinker, A., Friedman, D., Dugan, P., Winawer, J., & Groen, I. I. A. (2024). Temporal Dynamics of Short-Term Neural Adaptation across Human Visual Cortex. PLOS Computational Biology, 20(5), e1012161. 10.1371/journal.pcbi.1012161

Burlingham, C. S., Mirbagheri, S., & Heeger, D. J. (2022). A Unified Model of the Task-Evoked Pupil Response. Science Advances, 8(16), eabi9979. 10.1126/sciadv.abi9979

Chehab, O., Défossez, A., Jean-Christophe, L., Gramfort, A., & King, J.-R. (2022). Deep Recurrent Encoder: An End-to-End Network to Model Magnetoencephalography at Scale. Neurons, Behavior, Data Analysis, And Theory. 10.51628/001c.38668

Coco, M. I., Nuthmann, A., & Dimigen, O. (2020). Fixation-Related Brain Potentials during Semantic Integration of Object-Scene Information. Journal of Cognitive Neuroscience, 32(4), 571–589. 10.1162/jocn_a_01504

Cox, R. W., & Hyde, J. S. (1997). Software Tools for Analysis and Visualization of fMRI Data. NMR in Biomedicine, 10(4–5), 171–178. 10.1002/(SICI)1099-1492(199706/08)10:4/5<171::AID-NBM453>3.0.CO;2-L

Danisch, S., & Krumbiegel, J. (2021). Makie.jl: Flexible high-performance data visualization for Julia. Journal of Open Source Software, 6(65), 3349. 10.21105/joss.03349

Davis, H. (1936). ACTION POTENTIALS OF THE BRAIN: IN NORMAL PERSONS AND IN NORMAL STATES OF CEREBRAL ACTIVITY. Archives of Neurology & Psychiatry, 36(6), 1214. 10.1001/archneurpsyc.1936.02260120061004

Der, G., & Deary, I. J. (2006). Age and Sex Differences in Reaction Time in Adulthood: Results from the United Kingdom Health and Lifestyle Survey. Psychology and Aging, 21(1), 62–73. 10.1037/0882-7974.21.1.62

Dimigen, O., & Ehinger, B. V. (2021). Regression-Based Analysis of Combined EEG and Eye-Tracking Data: Theory and Applications. Journal of Vision, 21(1), 3. 10.1167/jov.21.1.3

Dimigen, O., Sommer, W., Hohlfeld, A., Jacobs, A. M., & Kliegl, R. (2014). Co-Registration of Eye Movements and EEG in Natural Reading: Analyses & Review. The Mind Research Repository (Beta), 0(1).

Ehinger, B. V., & Dimigen, O. (2019). Unfold: An Integrated Toolbox for Overlap Correction, Non-Linear Modeling, and Regression-Based EEG Analysis. Peerj, 7, e7838. 10.7717/peerj.7838

Ehinger, B. V., Kaufhold, L., & König, P. (2018). Probing the Temporal Dynamics of the Exploration–Exploitation Dilemma of Eye Movements. Journal of Vision, 18(3), 6. 10.1167/18.3.6

Ehinger, B., & Alday, P. (2024, September). Unfold.Jl: Event-Related Regression Toolbox. 10.5281/zenodo.10422370

Ehinger, B., & Alday, P. (2025, April 4). Unfold.Jl: Event-Related Regression Toolbox. 10.5281/zenodo.15145161

Ehinger, B., & Skukies, R. (2025, May 16). UnfoldBIDS. https://github.com/unfoldtoolbox/UnfoldBIDS.jl

Esteban, O., Markiewicz, C. J., Blair, R. W., Moodie, C. A., Isik, A. I., Erramuzpe, A., Kent, J. D., Goncalves, M., DuPre, E., Snyder, M., Oya, H., Ghosh, S. S., Wright, J., Durnez, J., Poldrack, R. A., & Gorgolewski, K. J. (2019). fMRIPrep: A Robust Preprocessing Pipeline for Functional MRI. Nature Methods, 16(1), 111–116. 10.1038/s41592-018-0235-4

Frömer, R., Maier, M., & Abdel Rahman, R. (2018). Group-Level EEG-Processing Pipeline for Flexible Single Trial-Based Analyses Including Linear Mixed Models. Frontiers in Neuroscience, 12. 10.3389/fnins.2018.00048

Gehring, J., Ignatiadis, N., & Alday, P. (2023, July). Juliangehring/MultipleTesting.Jl: MultipleTesting v0.6.0. 10.5281/zenodo.8110623

Gert, A. L., Ehinger, B. V., Timm, S., Kietzmann, T. C., & König, P. (2022). WildLab: A Naturalistic Free Viewing Experiment Reveals Previously Unknown Electroencephalography Signatures of Face Processing. European Journal of Neuroscience, 56(11), 6022–6038. 10.1111/ejn.15824

Giannoni, A., Baruah, R., Leong, T., Rehman, M. B., Pastormerlo, L. E., Harrell, F. E., Coats, A. J. S., & Francis, D. P. (2014). Do Optimal Prognostic Thresholds in Continuous Physiological Variables Really Exist? Analysis of Origin of Apparent Thresholds, with Systematic Review for Peak Oxygen Consumption, Ejection Fraction and BNP. Plos One, 9(1), e81699. 10.1371/journal.pone.0081699

Gilbert, C. D., & Sigman, M. (2007). Brain States: Top-Down Influences in Sensory Processing. Neuron, 54(5), 677–696. 10.1016/j.neuron.2007.05.019

Glover, G. H. (1999). Deconvolution of Impulse Response in Event-Related BOLD fMRI1. Neuroimage, 9(4), 416–429. 10.1006/nimg.1998.0419

Gramann, K., McKendrick, R., Baldwin, C., Roy, R. N., Jeunet, C., Mehta, R. K., & Vecchiato, G. (2021). Grand Field Challenges for Cognitive Neuroergonomics in the Coming Decade. Frontiers in Neuroergonomics, 2. 10.3389/fnrgo.2021.643969

Greenland, S. (2017). Invited Commentary: The Need for Cognitive Science in Methodology. American Journal of Epidemiology, 186(6), 639–645. 10.1093/aje/kwx259

Greve, D. N., & Fischl, B. (2009). Accurate and Robust Brain Image Alignment Using Boundary-Based Registration. Neuroimage, 48(1), 63–72. 10.1016/j.neuroimage.2009.06.060

Grinband, J., Wager, T. D., Lindquist, M. A., Ferrera, V. P., & Hirsch, J. (2008). Detection of Time-Varying Signals in Event-Related fMRI Designs. Neuroimage, 43(3), 509–520. 10.1016/j.neuroimage.2008.07.065

Groen, I. I. A., Piantoni, G., Montenegro, S., Flinker, A., Devore, S., Devinsky, O., Doyle, W., Dugan, P., Friedman, D., Ramsey, N. F., Petridou, N., & Winawer, J. (2022). Temporal Dynamics of Neural Responses in Human Visual Cortex. Journal of Neuroscience, 42(40), 7562–7580. 10.1523/JNEUROSCI.1812-21.2022

Hassall, C. D., Harley, J., Kolling, N., & Hunt, L. T. (2022). Temporal Scaling of Human Scalp-Recorded Potentials. Proceedings of the National Academy of Sciences, 119(43), e2214638119. 10.1073/pnas.2214638119

Hauk, O., Davis, M. H., Ford, M., Pulvermüller, F., & Marslen-Wilson, W. D. (2006). The Time Course of Visual Word Recognition as Revealed by Linear Regression Analysis of ERP Data. Neuroimage, 30(4), 1383–1400. 10.1016/j.neuroimage.2005.11.048

Hershman, R., & Henik, A. (2019). Dissociation between Reaction Time and Pupil Dilation in the Stroop Task. Journal of Experimental Psychology. Learning, Memory, And Cognition, 45(10), 1899–1909. 10.1037/xlm0000690

Isabella, S. L., Urbain, C., Cheyne, J. A., & Cheyne, D. (2019). Pupillary Responses and Reaction Times Index Different Cognitive Processes in a Combined Go/Switch Incidental Learning Task. Neuropsychologia, 127, 48–56. 10.1016/j.neuropsychologia.2019.02.007

Jenkinson, M., Bannister, P., Brady, M., & Smith, S. (2002). Improved Optimization for the Robust and Accurate Linear Registration and Motion Correction of Brain Images. Neuroimage, 17(2), 825–841. 10.1006/nimg.2002.1132

JuliaDSP/DSP.Jl. (2025, July 4). https://github.com/JuliaDSP/DSP.jl

JuliaNeuroscience/NIfTI.Jl. (2025, March 4). https://github.com/JuliaNeuroscience/NIfTI.jl

Jung, T. P., Makeig, S., Westerfield, M., Townsend, J., Courchesne, E., & Sejnowski, T. J. (1999). Independent Component Analysis of Single-Trial Event-Related Potentials.

Kappenman, E. S., Farrens, J. L., Zhang, W., Stewart, A. X., & Luck, S. J. (2021). ERP CORE: An Open Resource for Human Event-Related Potential Research. Neuroimage, 225, 117465. 10.1016/j.neuroimage.2020.117465

Kilavik, B. E., Confais, J., Ponce-Alvarez, A., Diesmann, M., & Riehle, A. (2010). Evoked Potentials in Motor Cortical Local Field Potentials Reflect Task Timing and Behavioral Performance. Journal of Neurophysiology, 104(5), 2338–2351. 10.1152/jn.00250.2010

Kristensen, E., Guerin-Dugué, A., & Rivet, B. (2017). Regularization and a General Linear Model for Event-Related Potential Estimation. Behavior Research Methods, 49(6), 2255–2274. 10.3758/s13428-017-0856-z

Lange, K., & Röder, B. (2006). Orienting Attention to Points in Time Improves Stimulus Processing Both within and across Modalities. Journal of Cognitive Neuroscience, 18(5), 715–729. 10.1162/jocn.2006.18.5.715

Larsen, R. S., & Waters, J. (2018). Neuromodulatory Correlates of Pupil Dilation. Frontiers in Neural Circuits, 12, 21. 10.3389/fncir.2018.00021

Luck, S. J. (2014). An Introduction to the Event-Related Potential Technique, Second Edition. MIT Press.

Luck, S. J. (2022). Applied Event-Related Potential Data Analysis. LibreTexts.

Mathewson, K. E., Kuziek, J. P., Scanlon, J. E. M., & Robles, D. (2024). The Moving Wave: Applications of the Mobile EEG Approach to Study Human Attention. Psychophysiology, 61(9), e14603. 10.1111/psyp.14603

Mikheev, V., Döring, S., Gärtner, N., Baumgartner, D., & Ehinger, B. (2024b, September). UnfoldMakie. 10.5281/zenodo.13747561

Mikheev, V., Skukies, R., & Ehinger, B. V. (2024a). The Art of Brainwaves: A Survey on Event-Related Potential Visualization Practices. Aperture Neuro, 4. 10.52294/001c.116386

Momenian, M., Vaghefi, M., Sadeghi, H., Momtazi, S., & Meyer, L. (2024). Language Prediction in Monolingual and Bilingual Speakers: An EEG Study. Scientific Reports, 14(1), 6818. 10.1038/s41598-024-57426-y

Mumford, J. A., Bissett, P. G., Jones, H. M., Shim, S., Rios, J. A. H., & Poldrack, R. A. (2023). The Response Time Paradox in Functional Magnetic Resonance Imaging Analyses. Nature Human Behaviour, 1–12. 10.1038/s41562-023-01760-0

Nakuci, J., Yeon, J., Kim, J.-H., Kim, S.-P., & Rahnev, D. (2024). Behavior Can Be Decoded across the Cortex When Individual Differences Are Considered. Imaging Neuroscience, 2, 1–17. 10.1162/imag_a_00359

Nikolaev, A. R., Ehinger, B. V., Meghanathan, R. N., & Van Leeuwen, C. (2023). Planning to Revisit: Neural Activity in Refixation Precursors. Journal of Vision, 23(7), 2. 10.1167/jov.23.7.2

Nikolaev, A. R., Meghanathan, R. N., & van Leeuwen, C. (2016). Combining EEG and Eye Movement Recording in Free Viewing: Pitfalls and Possibilities. Brain and Cognition, 107, 55–83. 10.1016/j.bandc.2016.06.004

Nuthmann, A. (2017). Fixation Durations in Scene Viewing: Modeling the Effects of Local Image Features, Oculomotor Parameters, and Task. Psychonomic Bulletin & Review, 24(2), 370–392. 10.3758/s13423-016-1124-4

Penny, W. D., Friston, K. J., Ashburner, J. T., Kiebel, S. J., & Nichols, T. E. (2011). Statistical Parametric Mapping: The Analysis of Functional Brain Images. Elsevier.

Pernet, C. R., Chauveau, N., Gaspar, C., & Rousselet, G. A. (2011). LIMO EEG: A Toolbox for Hierarchical LInear MOdeling of ElectroEncephaloGraphic Data. Computational Intelligence and Neuroscience, 2011, e831409. 10.1155/2011/831409

Philiastides, M. G., & Sajda, P. (2007). EEG-Informed fMRI Reveals Spatiotemporal Characteristics of Perceptual Decision Making. Journal of Neuroscience, 27(48), 13082–13091. 10.1523/JNEUROSCI.3540-07.2007

Poli, R., Cinel, C., Citi, L., & Sepulveda, F. (2010). Reaction-Time Binning: A Simple Method for Increasing the Resolving Power of ERP Averages. Psychophysiology, 47(3), 467–485. 10.1111/j.1469-8986.2009.00959.x

Pruim, R. H. R., Mennes, M., van Rooij, D., Llera, A., Buitelaar, J. K., & Beckmann, C. F. (2015). ICA-AROMA: A Robust ICA-based Strategy for Removing Motion Artifacts from fMRI Data. Neuroimage, 112, 267–277. 10.1016/j.neuroimage.2015.02.064

Qian, J., Hastie, T., Friedman, J., Tibshirani, R., & Simon, N. (2013,). Glmnet in Matlab.

Rousselet, G. A. (2025). Using Cluster-Based Permutation Tests to Estimate MEG/EEG Onsets: How Bad Is It?. European Journal of Neuroscience, 61(1), e16618. 10.1111/ejn.16618

Schaefer, A., Kong, R., Gordon, E., Laumann, T., Zuo, X., Holmes, A., Eickhoff, S., & Btt, Y. (2018). Local-Global Parcellation of the Human Cerebral Cortex from Intrinsic Functional Connectivity MRI. Cerebral Cortex (New York, N.Y. : 1991), 28(9). 10.1093/cercor/bhx179

Senn, S. (2005). Dichotomania: An Obsessive Compulsive Disorder That Is Badly Affecting the Quality of Analysis of Pharmaceutical Trials. International Statistical Istitute 55th Session, 13.

Skerritt-Davis, B., & Elhilali, M. (2018). Detecting Change in Stochastic Sound Sequences. PLOS Computational Biology, 14(5), e1006162. 10.1371/journal.pcbi.1006162

Skukies, R. S. (2020). Validation of Measures of Consciousness Using Propofol Anesthesia.

Smith, N. J., & Kutas, M. (2015b). Regression-Based Estimation of ERP Waveforms: I. The rERP Framework. Psychophysiology, 52(2), 157–168. 10.1111/psyp.12317

Smith, N. J., & Kutas, M. (2015a). Regression-Based Estimation of ERP Waveforms: II. Non-linear Effects, Overlap Correction, and Practical Considerations. Psychophysiology, 52(2), 169–181. 10.1111/psyp.12320

Snowden, R. J., O’Farrell, K. R., Burley, D., Erichsen, J. T., Newton, N. V., & Gray, N. S. (2016). The Pupil’s Response to Affective Pictures: Role of Image Duration, Habituation, and Viewing Mode. Psychophysiology, 53(8), 1217–1223. 10.1111/psyp.12668

Strauch, C., Koniakowsky, I., & Huckauf, A. (2020). Decision Making and Oddball Effects on Pupil Size: Evidence for a Sequential Process. 3(1), 7. 10.5334/joc.96

Stroop, J. R. (1935). Studies of Interference in Serial Verbal Reactions. Journal of Experimental Psychology, 18(6), 643–662. 10.1037/h0054651

Sun, J., Osth, A. F., & Feuerriegel, D. (2024). The Late Positive Event-Related Potential Component Is Time Locked to the Decision in Recognition Memory Tasks. Cortex. 10.1016/j.cortex.2024.04.017

Van Humbeeck, N., Meghanathan, R. N., Wagemans, J., van Leeuwen, C., & Nikolaev, A. R. (2018). Presaccadic EEG Activity Predicts Visual Saliency in Free-Viewing Contour Integration. Psychophysiology, 55(12), e13267. 10.1111/psyp.13267

Wang, J., Narain, D., Hosseini, E. A., & Jazayeri, M. (2018). Flexible Timing by Temporal Scaling of Cortical Responses. Nature Neuroscience, 21(1), 102–110. 10.1038/s41593-017-0028-6

Welke, D., & Vessel, E. A. (2022, April). Naturalistic Viewing Conditions Can Increase Task Engagement and Aesthetic Preference but Have Only Minimal Impact on EEG Quality (p. 2021). bioRxiv. 10.1101/2021.09.18.460905

Winkler, A. M., Taylor, P. A., Nichols, T. E., & Rorden, C. (2024, January 7). False Discovery Rate and Localizing Power. 10.48550/arXiv.2401.03554

Wood, S. N. (2017). Generalized Additive Models: An Introduction with R, Second Edition (2nd ed.). Chapman and Hall/CRC. 10.1201/9781315370279

Yarkoni, T., Barch, D. M., Gray, J. R., Conturo, T. E., & Braver, T. S. (2009). BOLD Correlates of Trial-by-Trial Reaction Time Variability in Gray and White Matter: A Multi-Study fMRI Analysis. PLOS ONE, 4(1), e4257. 10.1371/journal.pone.0004257

