## Appendix for "Brain responses vary in duration - modeling strategies and challenges"

### APPENDIX A

#### A.1 Noise Preprocessing

Eyes closed resting state data was preprocessed as follows

1. Data was down-sampled to 500Hz
2. EEGLAB's `pop_clean_rawdata()` was used to reject bad channels using Parameters: `FlatlineCriterion = 5`; `ChannelCriterion = 0.8`; `LineNoiseCriterion = 4`; `Highpass = [0.25, 0.75]`
3. Data was re-referenced to average reference
4. A copy of the data was made, and the copy was high-pass filtered at 1.5Hz
5. ICA was calculated on the (filtered) copied data
6. Bad components were marked using ICLabel with Parameters only set for eye movements and muscle artefacts (i.e. only eye- and muscle-artefacts were rejected)
7. ICA weights and reject-markers were copied to original data and bad components were subtracted
8. Rejected channels were interpolated via spherical interpolation

Additionally, during simulations the data was down-sampled (default 100Hz) and high-pass filtered (default 0.5 Hz) to match the simulations.

#### A.2 No Noise Results

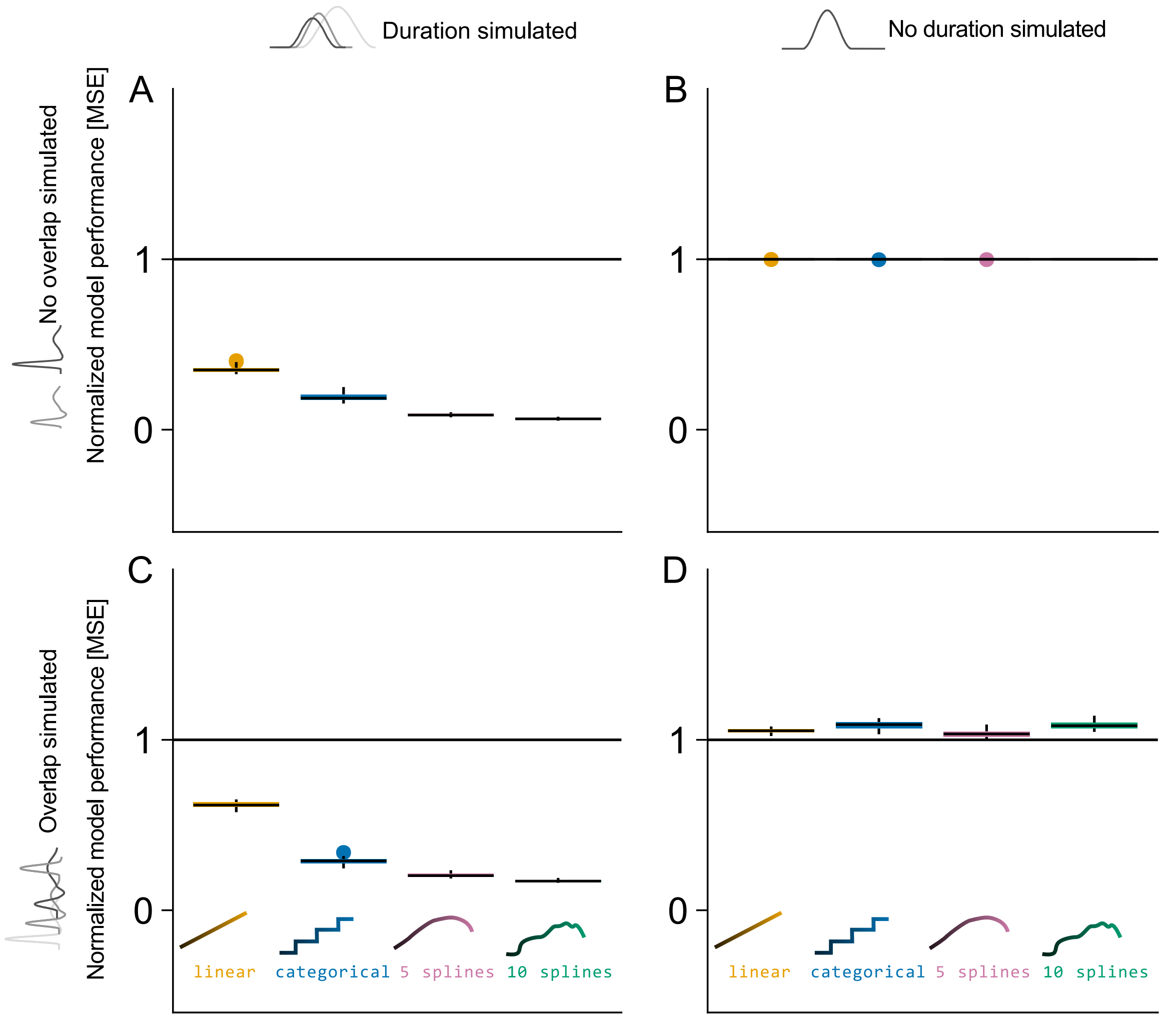


Figure 10: Normalized mean squared error results for the four tested models (including duration as: linear, categorical, 5-spline, 10-spline variable) in different simulation settings. Black line (y-value one) indicated results from classical averaging; MSE of zero indicated perfect estimation of the ground truth. (A) Results when a duration effect, but no overlap is simulated. The spline strategies outperform the other strategies. (B) Results when no duration effect and no overlap were simulated, but duration effects were still estimated. Little overfit is visible here. (C) Results when duration effects were simulated, signals overlap, and overlap correction was used for modeling. No interaction between duration modeling and overlap correction was observed on the MSE performance. (D) Results when duration effect was not simulated, and signals overlap, and results are overlap-corrected.

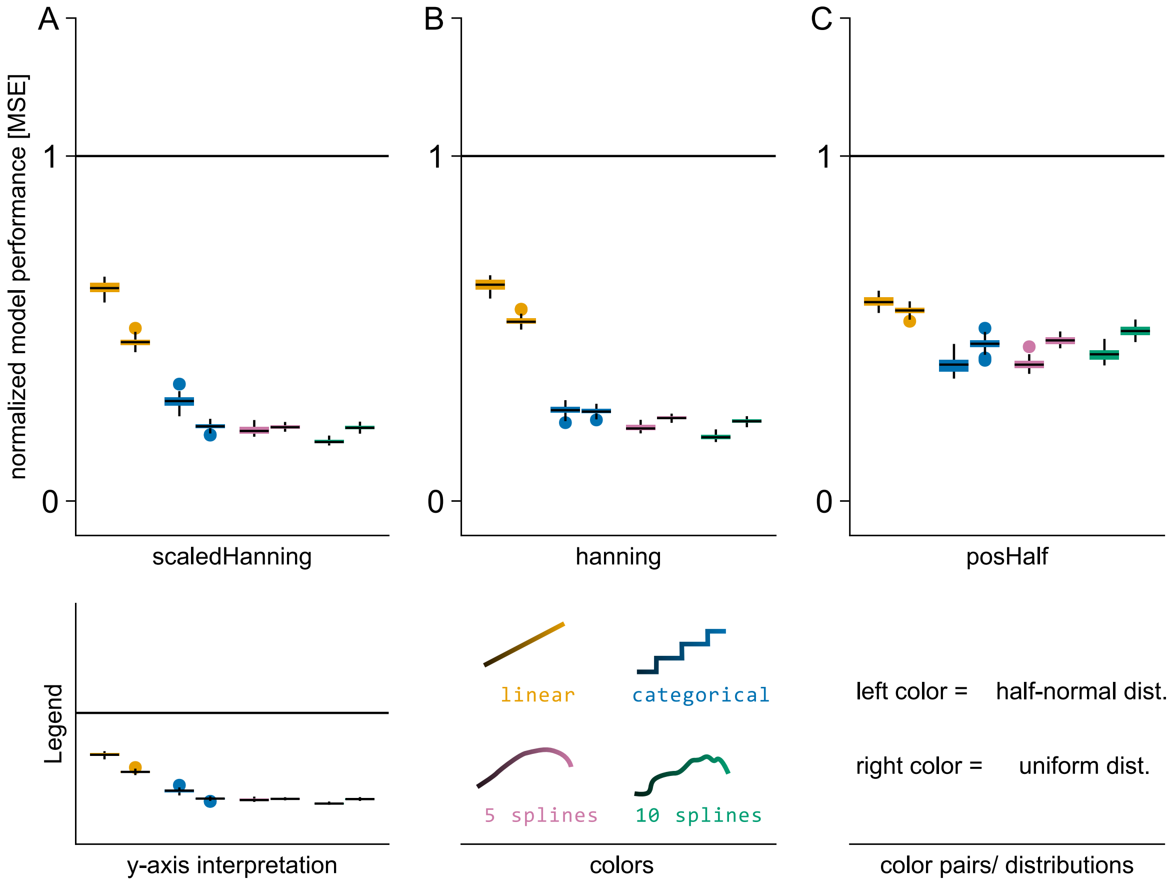


Figure 11: Normalized mean squared error results for the four tested models in simulations without noise (including duration as: linear, categorical, 5-spline, 10-spline variable) between shapes (A-C) and duration distributions (color pairs). Black line (y-value one) indicated results from classical averaging; MSE of zero indicated perfect estimation of the ground truth. Parameter settings for all panels: duration affects shape; overlap simulated; overlap-corrected/ modeled.

#### A.3 Duration Modeling vs. Overlap Correction

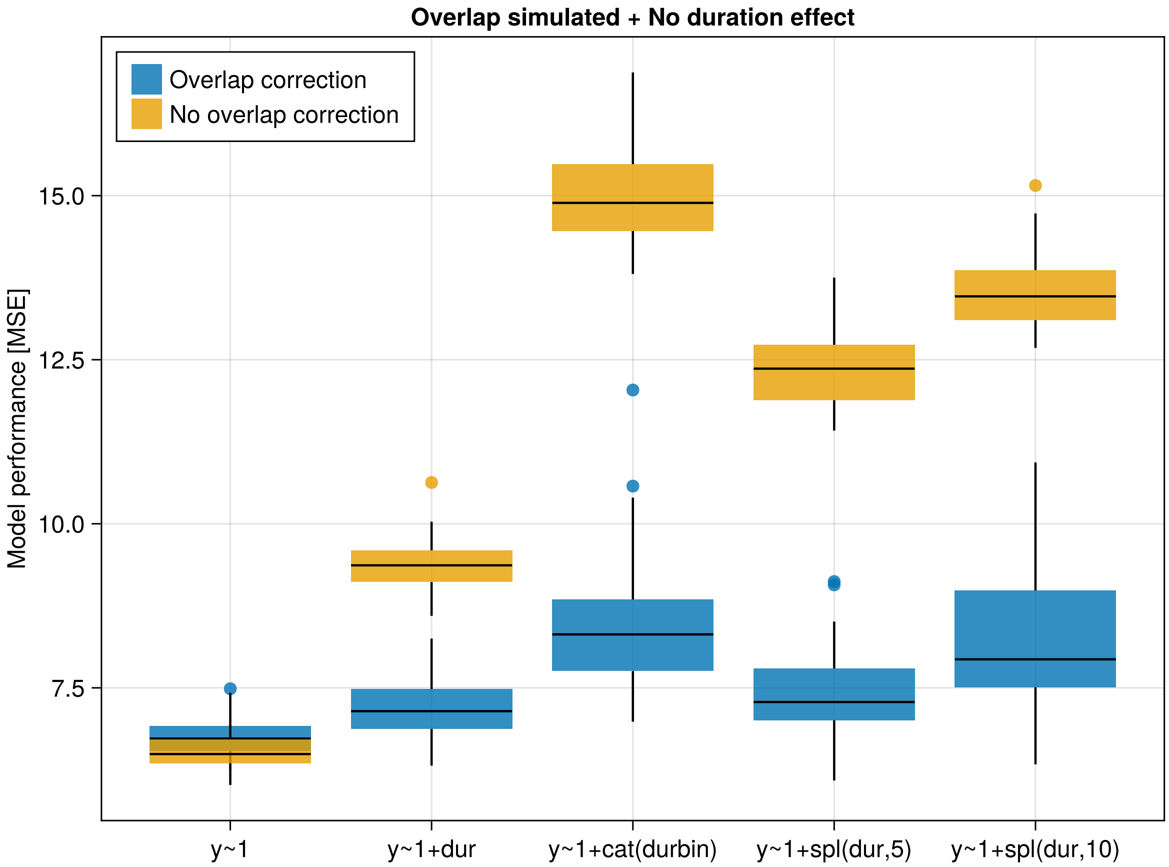


Figure 12: *Not* normalized mean square error results when overlap is simulated, but no duration effect is simulated for five different models (classical averaging ($y \sim 1$) plus the four tested models (duration as: linear, categorical, 5-spline, 10-spline variable)), once with overlap correction (blue) and once without overlap correction (orange).

#### A.4 MSE Plots of Baseline Corrected Data

Baseline was corrected for each marginalized ERP independently in with a baseline period of −0.5s to 0.0s.

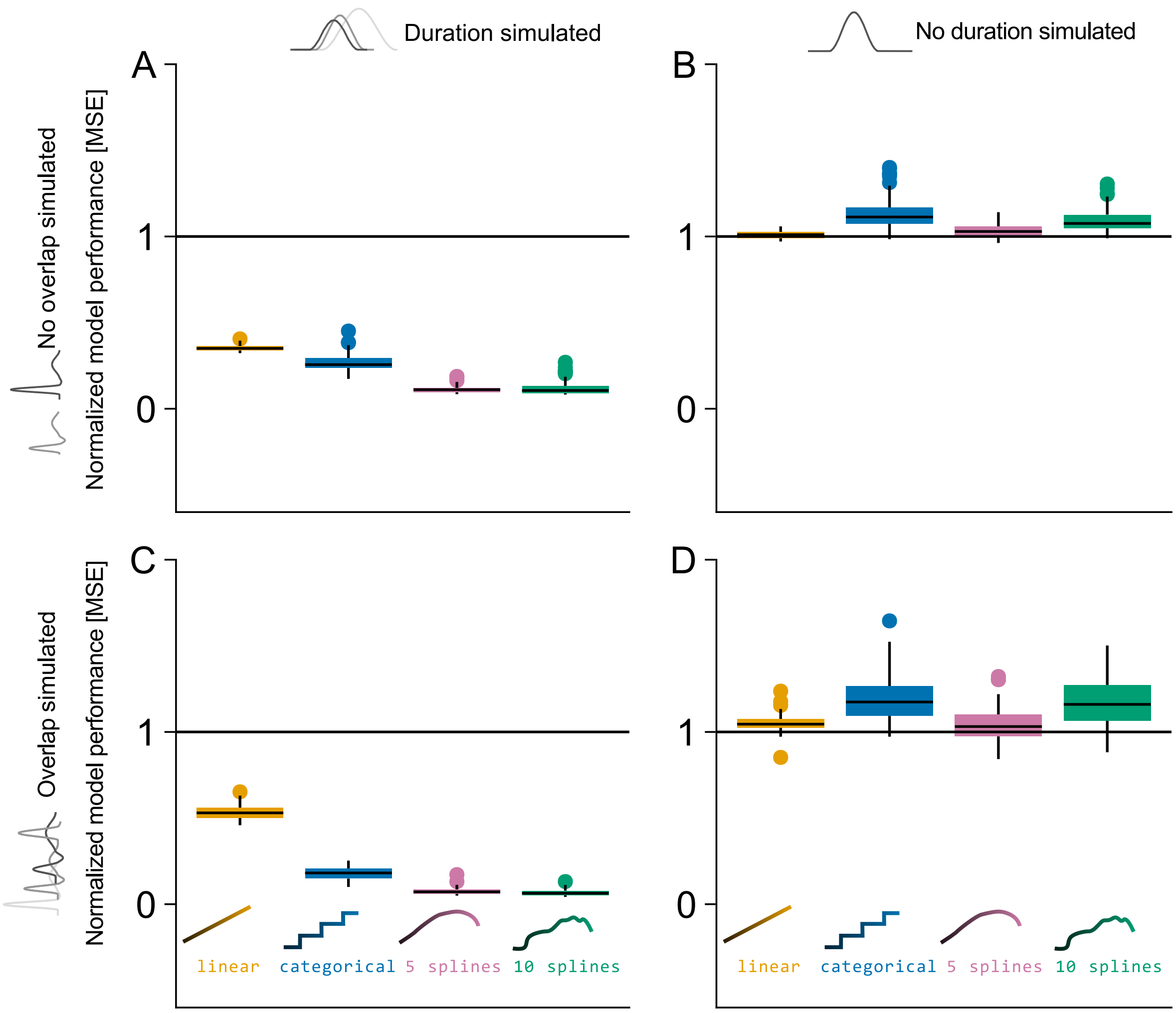


Figure 13: Normalized mean squared error results of baseline corrected estimates for the four tested models (including duration as: linear, categorical, 5-spline, 10-spline variable) in different simulation settings. Black line (y-value one) indicated results from classical averaging; MSE of zero indicated perfect estimation of the ground truth. (A) Results when a duration effect, but no overlap is simulated. The spline strategies outperform the other strategies. (B) Results when no duration effect and no overlap were simulated, but duration effects were still estimated. Little overfit is visible here. (C) Results when duration effects were simulated, signals overlap, and overlap correction was used for modeling. No interaction between duration modeling and overlap correction was observed on the MSE performance. (D) Results when duration effect was not simulated, and signals overlap, and results are overlap-corrected.

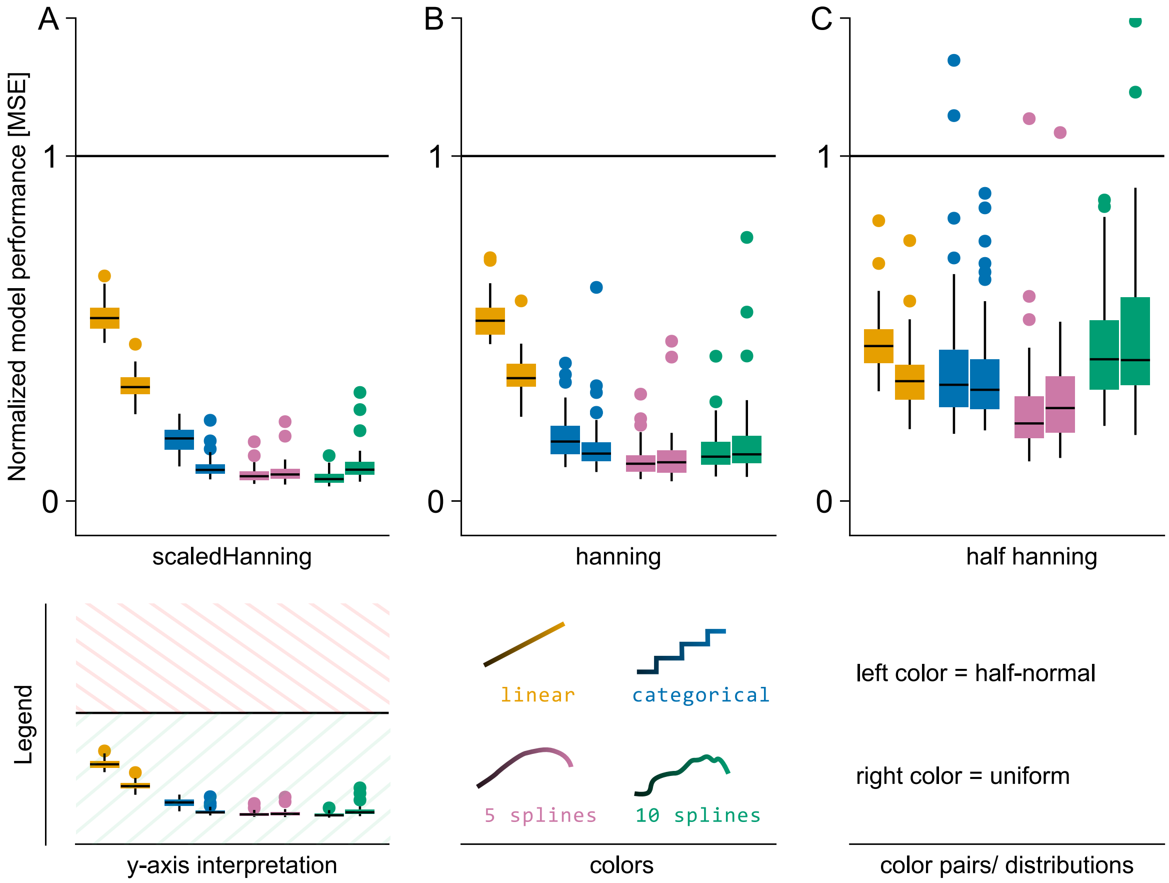


Figure 14: Normalized mean squared error results of baseline corrected estimates for the four tested models in simulations without noise (including duration as: linear, categorical, 5-spline, 10-spline variable) between shapes (A-C) and duration distributions (color pairs). Black line (y-value one) indicated results from classical averaging; MSE of zero indicated perfect estimation of the ground truth. Parameter settings for all panels: duration affects shape; overlap simulated; overlap-corrected/ modeled.

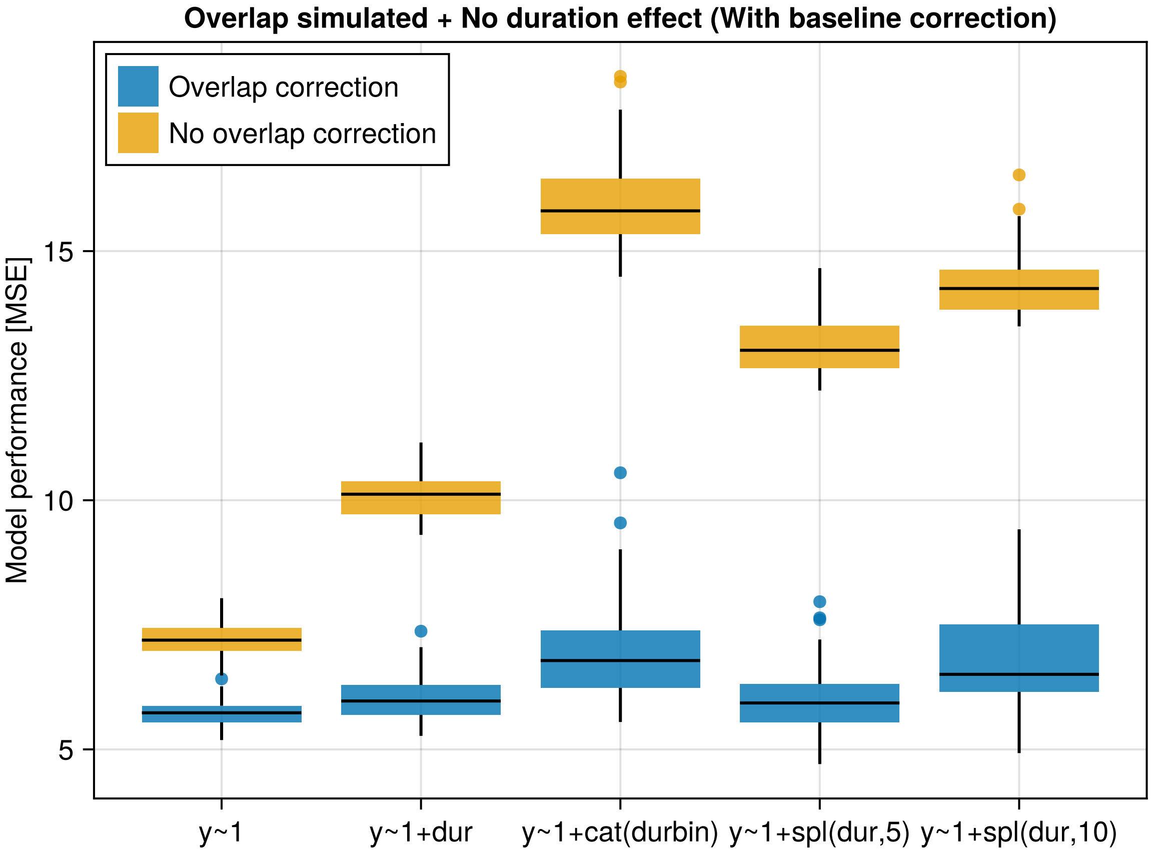


Figure 15: *Not* normalized mean square error results of baseline corrected estimates when overlap is simulated, but no duration effect is simulated for five different models (classical averaging ($y \sim 1$) plus the four tested models (duration as: linear, categorical, 5-spline, 10-spline variable)), once with overlap correction (blue) and once without overlap correction (orange).

#### A.5 HRF Parameters

- p(1) - delay of response (relative to onset): 6
- p(2) - delay of undershoot (relative to onset): 16
- p(3) - dispersion of response: 1
- p(4) - dispersion of undershoot: 1
- p(5) - ratio of response to undershoot: 6
- p(6) - onset {seconds}: 0
- p(7) - length of kernel {seconds}: 32
